# Long-term learning transforms prefrontal cortex representations during working memory

**DOI:** 10.1101/2022.02.22.481537

**Authors:** Jacob A. Miller, Arielle Tambini, Anastasia Kiyonaga, Mark D’Esposito

## Abstract

The lateral prefrontal cortex (lPFC) is reliably active during working memory (WM) across human and animal models, but the role of lPFC in successful WM is under debate. For instance, non-human primate (NHP) electrophysiology research finds that lPFC circuitry stores WM representations. Human neuroimaging instead suggests that lPFC plays a control function over WM content that is stored in sensory cortices. These seemingly incompatible WM accounts are often confounded by differences in the amount of task training and stimulus exposure across studies (i.e., NHPs tend to be trained extensively). Here, we test the possibility that such long-term training may alter the role of lPFC in WM maintenance. We densely sampled WM-related activity across learning, in three human participants, using a longitudinal functional MRI (fMRI) protocol. Over three months, participants trained on (1) a serial reaction time (SRT) task, wherein complex fractal stimuli were embedded within probabilistic sequences, and (2) a delayed recognition task probing WM for trained or novel stimuli. Participants were scanned frequently throughout training, to track how WM activity patterns change with repeated stimulus exposure and long-term associative learning. WM task performance improved for trained (but not novel) fractals and, neurally, delay activity significantly increased in distributed lPFC voxels across learning. Pattern similarity analyses also found that item-level WM representations became detectable within lPFC, but not in sensory cortices, and lPFC delay activity increasingly reflected sequence relationships from the SRT task, even though that information was task-irrelevant for WM. These findings demonstrate that human lPFC can show stimulus-selective WM responses with learning and WM representations are shaped by long-term experience. Therefore, influences from training and long-term memory may reconcile competing accounts of lPFC function during WM.

## Introduction

The lateral prefrontal cortex (lPFC) is considered critical for working memory (WM) across human and animal models (Funahashi et al., 1989; Goldman-Rakic, 1995; Leavitt et al., 2017; E. K. Miller et al., 2018; Sreenivasan et al., 2014). However, there is ongoing debate regarding the specific role that lPFC activity plays in successful WM (Christophel et al., 2017; Curtis & Sprague, 2021; Lara & Wallis, 2015; Mackey et al., 2016). Non-human primate (NHP) electrophysiology research typically finds that lPFC maintains feature-specific WM content (Constantinidis et al., 2018; Funahashi et al., 1989; Fuster & Alexander, 1971; Goldman-Rakic, 1995; E. K. Miller et al., 2018; Romo et al., 1999). Human neuroimaging suggests lPFC activity serves control functions over WM while feature-specific content is stored in sensory cortices instead (D’Esposito & Postle, 2015; Eriksson et al., 2015; Harrison & Tong, 2009; Riggall & Postle, 2012; Serences, 2016). However, these seemingly incompatible accounts are confounded by differences in species, measurement granularity, and the amount of task training that typically occurs across studies.

One possibility is that different indices of neural activity, across measurement scales, may support distinct conclusions about the cortical substrates for WM. That is, NHP studies typically record finer resolution single-unit neuronal activity compared to the millimeter scale of Blood Oxygen Level Dependent functional MRI (BOLD fMRI) (Mukamel et al., 2005; Park et al., 2017). Discrepancies between study findings may emerge if stimulus-specific WM content is represented in human lPFC, but undetectable at the coarser resolution of BOLD fMRI – for instance, via spiking patterns across populations of neurons that are spatially intermixed. The organization and spread of activity in sensory areas better matches the spatial resolution of BOLD fMRI, which may also reflect local field potentials from top-down modulation, in the absence of local spiking (Leavitt et al., 2017; Lorenc & Sreenivasan, 2021; Mendoza-Halliday et al., 2014; Serences, 2016). However, in some cases, stimulus-specific WM delay activity has been detected in human frontal cortex (Ester et al., 2015; Lee et al., 2013) or NHP sensory regions (Mendoza-Halliday et al., 2014; Supèr et al., 2001), highlighting the need to identify which factors truly drive observed differences in findings across studies.

In addition to differences in recording techniques between human and NHP studies, NHPs typically undergo months of training and perform orders of magnitude more task trials before the critical neural recordings occur (Berger et al., 2018; Birman & Gardner, 2016; Sarma et al., 2016). Humans typically complete only a few minutes of task practice prior to fMRI scanning. Differences observed in neural WM substrates across species may therefore be driven by long-term learning influences from extensive task and stimulus experience. In fact, the few studies that recorded from NHPs before and after WM training found plasticity in the form of increases in the magnitude of WM delay activity and in the strength of item-level stimulus representations in anterior lPFC (Dang et al., 2021; Meyer et al., 2011; Qi et al., 2019; Riley et al., 2018; Sarma et al., 2016). Human lPFC may likewise represent item-level information in WM depending on the level of prior training. However, the typical timeline of fMRI research has limited our ability to directly test this hypothesis that WM representations change with long-term learning.

The brain regions and neural mechanisms for WM are classically considered separate from long-term memory (LTM) systems (Squire & Zola-Morgan, 1991; Warrington & Shallice, 1969; Wickelgren, 1969). However, some WM theories predict that learned associations or semantic links between items should be reflected during WM maintenance (LaRocque et al., 2014; Oberauer, 2009), and growing evidence suggests a common neural machinery between WM and LTM (Beukers et al., 2021; Borders et al., 2021; Fukuda & Woodman, 2017; Hoskin et al., 2019; Lewis-Peacock & Norman, 2014; Nee & Jonides, 2011; Ranganath et al., 2003; Ranganath & Blumenfeld, 2005; Yonelinas, 2013). In some cases WM capacity is also greater for stimuli with extensive exposure and familiarity (Asp et al., 2021; Brady et al., 2016; Jackson & Raymond, 2008; Xie & Zhang, 2017), suggesting that WM and supporting neural mechanisms may change with stimulus experience.

Here, we examined the possibility that long-term learning transforms human lPFC WM activity. We asked whether stimulus selectivity emerges in human lPFC as a function of training, akin to the stimulus-specific WM activity patterns typically found in NHP studies. To do so, three human participants each completed over 20 sessions of whole-brain fMRI along with extensive at-home training across three months. During this time, participants repeatedly performed a delayed recognition WM task and a sequence learning task, which both employed a set of 18 novel fractal stimuli that were unique to each participant. First, we asked whether lPFC delay period WM activity changed in magnitude across learning. Widespread decreases in lPFC activity could suggest more automatic task processing with training. Activity increases, however, could suggest greater selectivity for the repeated task structure or individual WM stimuli, as persistent activity in WM is associated with stimulus-selective patterns (Constantinidis et al., 2018; Curtis & Sprague, 2021; Murray et al., 2017). We then tested whether representations of individual stimuli or associative structures emerged in multivariate WM activity patterns over the course of learning. If item-level lPFC activity patterns develop over time, it would suggest that differences in participant training may explain discrepant accounts of lPFC as either a source of control over WM (from human studies) versus the storage site for WM content (from single-unit NHP studies). Alternatively, long-term learning may enhance sensory cortex representations of WM content but induce no changes in lPFC, suggesting that differences in lPFC vs sensory-based WM storage models are driven by other factors than long-term learning. Finally, to understand how WM representations are shaped by associative learning, we asked if representations of associations between stimuli in shared temporal sequences (learned outside of the WM task) were reflected within WM activity patterns. To preview the results, long-term learning changed the distribution and stimulus information content of lPFC WM delay activity, indicating that WM maintenance mechanisms may be flexible to the extent and nature of prior experience with the WM information. These results suggest that differences in the extent of training across species may masquerade as differences in lPFC function.

## Results

### Intensive training improves WM performance for trained, but not novel, stimuli

To determine how long-term learning influences cortical activity patterns underlying WM maintenance, we trained three human participants on a set of fractal stimuli that was unique to each participant (**Figure 1a**) over three months. These stimuli had no pre-existing meaning and have been used to characterize the influence of long-term associative learning on neural selectivity (Ghazizadeh et al., 2018; Kim et al., 2015; Sakai & Miyashita, 1991). These complex stimuli were chosen to extend the timecourse of learning, and to necessitate a detailed item representation for successful task performance. During the study period, each participant completed approximately 24 scanning (fMRI) sessions along with at-home behavioral training sessions multiple times per week (**Figure 1b**). Here we analyze the first 17 fMRI sessions (∼13 weeks), after which point new fractals were added into the stimulus set for a second phase of the study. During each fMRI session, participants performed two primary tasks, a serial reaction time (SRT) task followed by a WM task (**Figure 1c-d**). The WM task entailed a single-item delayed recognition test wherein the WM sample was either a fractal stimulus from the training set or a novel fractal that appeared only during that session. Before the study began, participants completed one block (24 trials) of WM task practice with pilot stimuli that never appeared in the main experiment. The first time each participant saw their unique set of 18 training stimuli was during the first scanning session. The SRT task used the same 18 trained fractal stimuli, and 12 of the stimuli were embedded in high probability sequences (**Figure 1c**). The sequences were not directly related to the goals of the WM task (which was always to remember a single item), but we took advantage of the sequence structure to analyze whether item-level WM representations reflected associations (sequence-level and categorical) from the SRT task.

**Figure 1.**
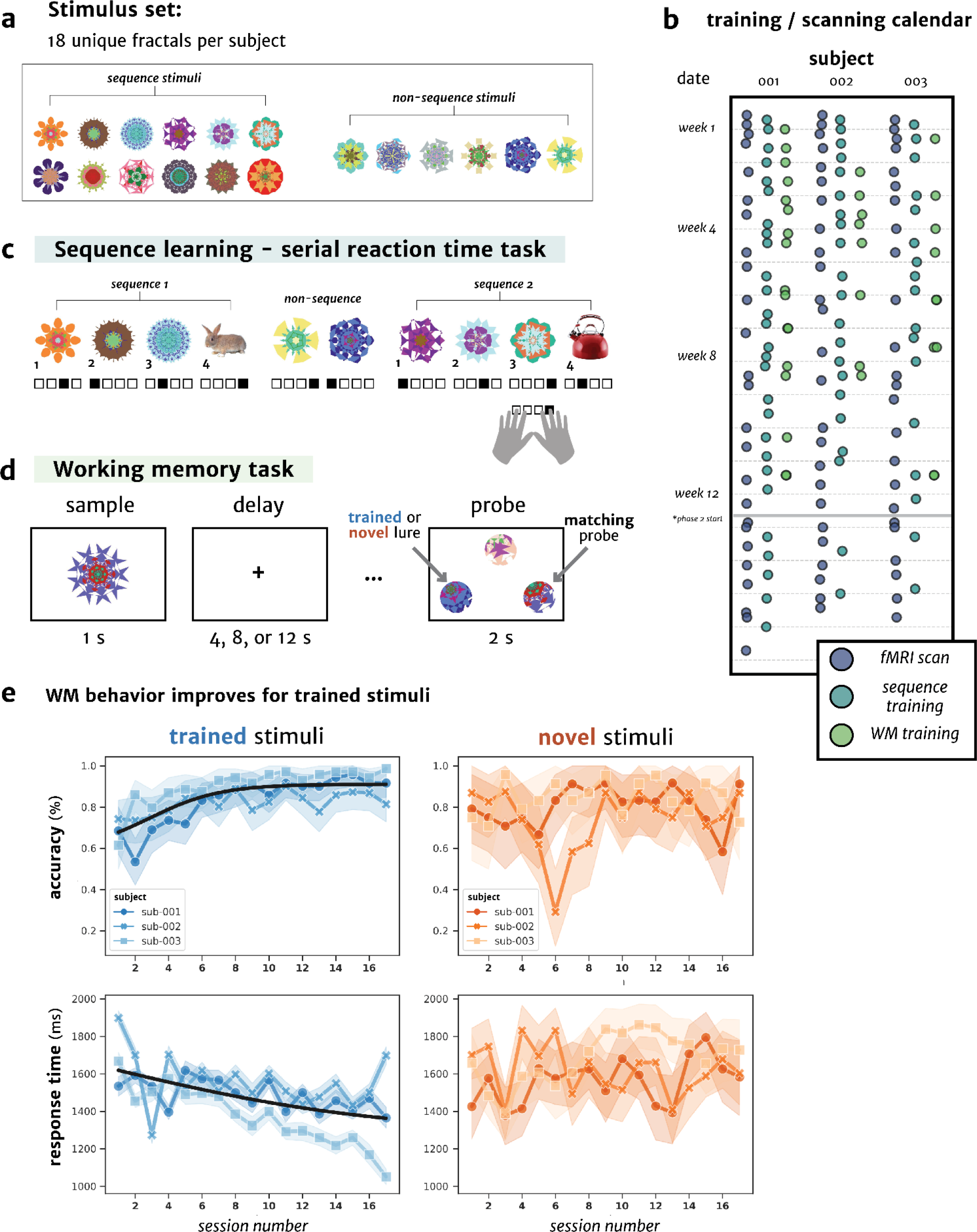
Longitudinal training across three months within individuals. **(a)** Example set of 18 unique fractal stimuli assigned to a single participant for the in-scanner and at-home behavioral tasks. **(b)** Calendar of all of the MRI (purple) and at-home sessions (SRT - dark green, WM - light green) for each of the three participants over the four months of the study. During each MRI session, participants completed both the sequence learning and WM tasks. The at-home training sessions consisted of modified versions of each task (**Methods**). The present study analyzes the first 17 sessions, as afterwards new stimuli were added into the training set for each participant. **(c)** The serial reaction time (SRT) task, in which each of the 18 trained stimuli was associated with one of four button responses. Of the 18 trained stimuli, 12 were part of 4 sequences that occurred with high probability (75%) in the SRT task, and participants learned the sequences over time (**SI Figure 1**). **(d)** The delayed three-alternative forced choice WM task, in which one fractal (trained or novel) was presented on each trial. After a jittered delay, participants indicated which occluded image matched the original sample. **(e)** WM task accuracy (top) and response time (bottom) improved across training (sessions 1-17) for trials with one of the 18 trained stimuli (*blue*), but not for trials with novel fractal stimuli (*orange*). Accuracy error bars represent a bootstrapped 68% confidence interval (C.I.) across different blocks within each session (4 per participant per session), while for RT, error bars (bootstrapped 68% C.I.) are plotted across trials within each session for each participant.

Across the course of training, behavior in the WM task improved for trained stimuli, but not for novel stimuli (**Figure 1e**). Mean WM probe accuracy (% correct) for trials with trained stimuli improved by 23% across the 17 sessions, whereas accuracy increased by 4% for novel stimuli. To characterize the change in WM performance over time, we used a fixed-effects, logistic modeling approach that can flexibly detect changes in learning over time, estimate *when* these changes are most prominent (inflection point), and adapt to different rates of learning (slopes; see **Figure 2a**). Significance in any change over time was assessed by correlating the predicted logistic model values with the actual data using cross-validation (**STAR Methods**: *Statistical methods*; see **Figure 2a** for schematic). There was a significant increase in WM accuracy for trained stimuli over time (*r* = 0.77, p < 0.001), and no reliable change for novel stimuli (*r* = 0.17, p = 0.07). This increase in WM accuracy for trained versus novel stimuli was confirmed by testing for an interaction between session number (1 → 17; mean-centered) and stimulus category (*trained* vs. *novel*) with a fixed-effects linear model (*t*(96) = 2.76, *p* = 0.007).

**Figure 2.**
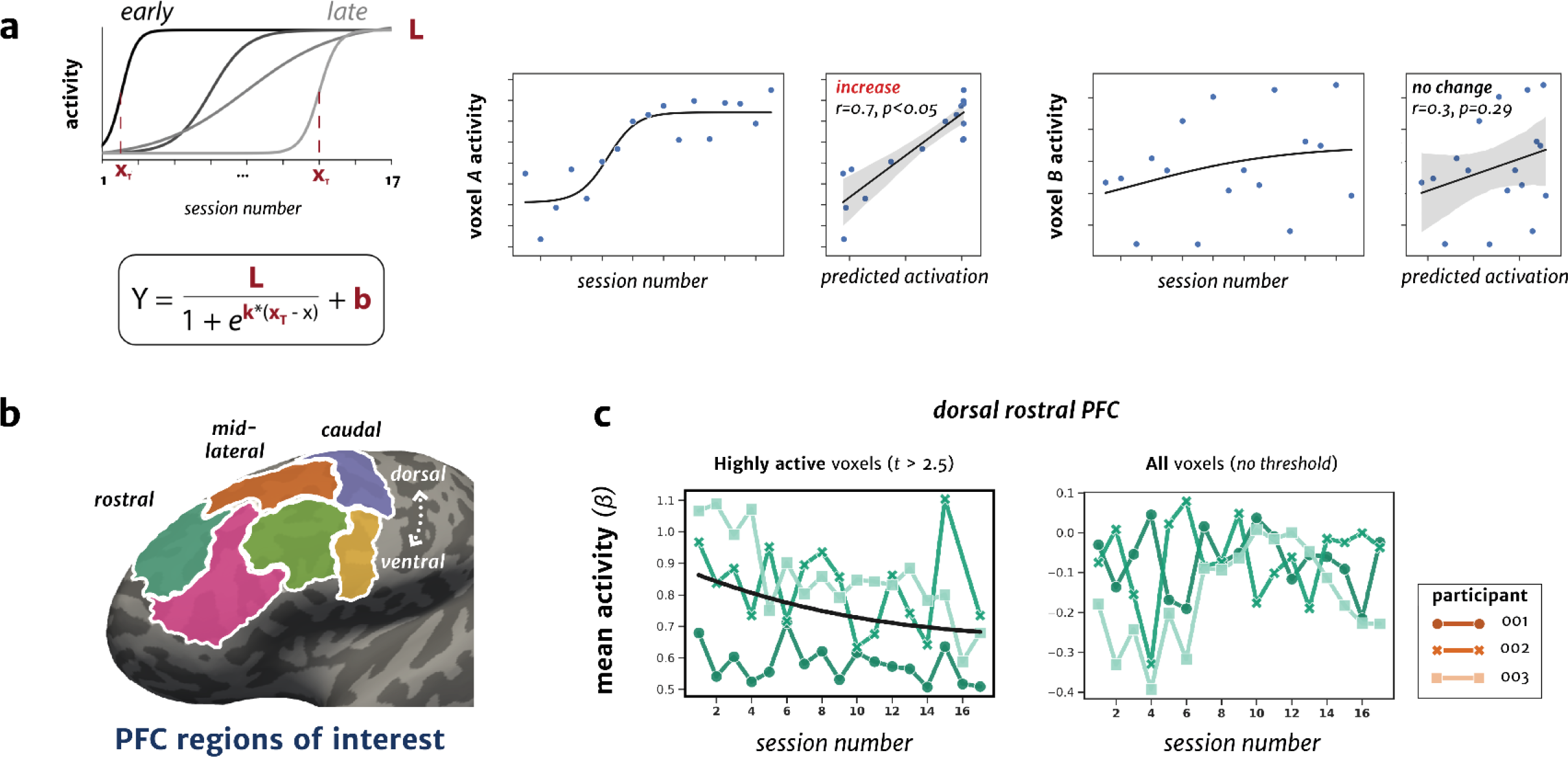
Mean WM delay activity changes in PFC across the course of learning. **(a)** Schematic of the logistic modeling approach, in which four free model parameters were fit to the mean WM delay activity from each voxel across sessions (**STAR Methods**). Left inset, example WM delay activity profiles from two voxels across sessions. Right inset, voxel activity correlated with the predicted activation from the logistic model after cross-validation. Positive r-values indicate increases in activity across sessions. **(b)** Six-region parcellation of the lateral PFC in an example participant’s inflated left hemisphere. The lPFC was divided along a rostral-caudal and dorsal-ventral axis by combining smaller parcels from a multi-modal atlas of the cerebral cortex (Glasser et al., 2016). The parcellation was designed to be homologous to NHP electrophysiology studies (Riley et al., 2018), and guided by functional subdivisions of human lPFC (Badre & D’Esposito, 2009). **(c)** Left: Mean activity for each fMRI session during the WM delay period for reliably active voxels (within each session), thresholded at *t* > 2.5. The dorsal rostral PFC ROI (*green*) showed a mean decrease in WM delay activity across sessions. Right: Mean activity for all voxels (unthresholded). For visualization, all ROIs with significant logistic model fits after FDR correction are indicated with a bolded plot border, along with the fitted logistic curve across sessions. No other ROIs showed a mean change in WM delay activity over the course of training.

A complementary pattern emerged when modeling WM probe response time (RT). For RT, there was also a significant interaction between session number and stimulus category (*t*(96) = -4.4, *p* < 0.001), which was driven by faster responses for trained stimuli over time (*r* = -0.54, *p* < 0.001), with no significant change for novel stimuli (*r* = 0.22, *p* = 0.06). The subsequent analyses use fixed-effects, logistic models, because they allow us to flexibly detect changes that occur at different times and rates across the 17 sessions (see **Figure 2a** for schematic), but all results generalize to a linear framework. Grouping data across participants with fixed-effects is also recommended for studies with a comparable sample size (Fries & Maris, 2021).

In parallel to the WM task, participants also learned associations between individual stimuli as part of regularly occurring sequences in the SRT task. Reliable associative learning across training was shown by reduced response times for intact sequences in the SRT task for all participants (**SI Figure 1**).

### Divergent changes in mean WM delay activity within dorsal PFC

To determine if lPFC activity changes across learning, we split the lPFC into six bilateral regions of interest (ROIs) along rostral-caudal (from the *frontal pole* to *precentral gyrus)* and dorsal-ventral (from the *superior frontal gyrus* to *inferior frontal gyrus*) axes (**Figure 2b**). This six-region parcellation was chosen to be homologous to a recent NHP study that recorded from multiple lPFC areas before and after WM training (Riley et al., 2018). We first tested for evidence of broad changes in WM delay activity over time. To test for changes in mean activity across entire ROIs, we considered two groups of voxels within each ROI. First, we examined whether peak activation in the WM delay period changed across sessions, which may reflect classical persistent activity during WM (Curtis & Sprague, 2021). To do this, we thresholded WM delay activity maps (collapsed across all delay lengths) for each participant and session at *t* > 2.5 and determined whether peak activation levels (*beta*-coefficient) changed over training (**Figure 2c**, *left*). Second, we analyzed the mean activity of all voxels across each ROI, without any thresholding, to ask whether there are changes across an entire cortical region (including voxels with lower WM activity). Changes in highly active voxels are sensitive to the magnitude of peak activity, but the precise location of highly activated voxels can shift from session to session.

The magnitude of WM delay activity changed across training in one lPFC area. That is, the peak WM delay period activity in dorsal rostral PFC decreased across sessions (*r* = -0.35, *p* = 0.007 [FDR-corrected *p* = 0.039]; **Figure 2c**, *left*), whereas the mean activity for all voxels in this area did not significantly change over sessions (*r* = 0.28, *p* = 0.02 [FDR-corrected *p* = 0.13], this model failed to converge with cross-validation, **Figure 2c**, *right*). No other ROIs showed training-related changes in either the peak WM delay activity or mean across all voxels (FDR-corrected *p-values* > 0.1; **SI Figure 2**). However, this approach may obscure divergent changes that occur within specific populations of voxels with learning. We next used a voxelwise regression approach to directly test whether individual voxels increased or decreased their activity over time.

### More cortical territory in PFC is recruited for WM delay activity across learning

Populations of voxels involved in WM maintenance may change their activity over training, as the stimuli and task become increasingly well-learned. For example, WM processing could become more “efficient” by recruiting less cortical territory. Or, more cortical territory could be engaged in representing and processing newly learned stimuli and task dimensions. To test these different predictions, for each voxel, we assessed the relationship between WM delay activity and training session with a logistic modeling approach using cross-validation (**Figure 2a**). We tested whether a meaningful proportion of individual voxels within each frontal ROI show systematic changes in activity over training compared to chance (permutation testing, see **STAR Methods**). A schematic of this voxelwise approach is shown in **Figure 3a**, allowing us to test whether populations of voxels in each ROI show divergent increases or decreases in WM delay activity with training—information that would be lost when averaging across voxels.

**Figure 3.**
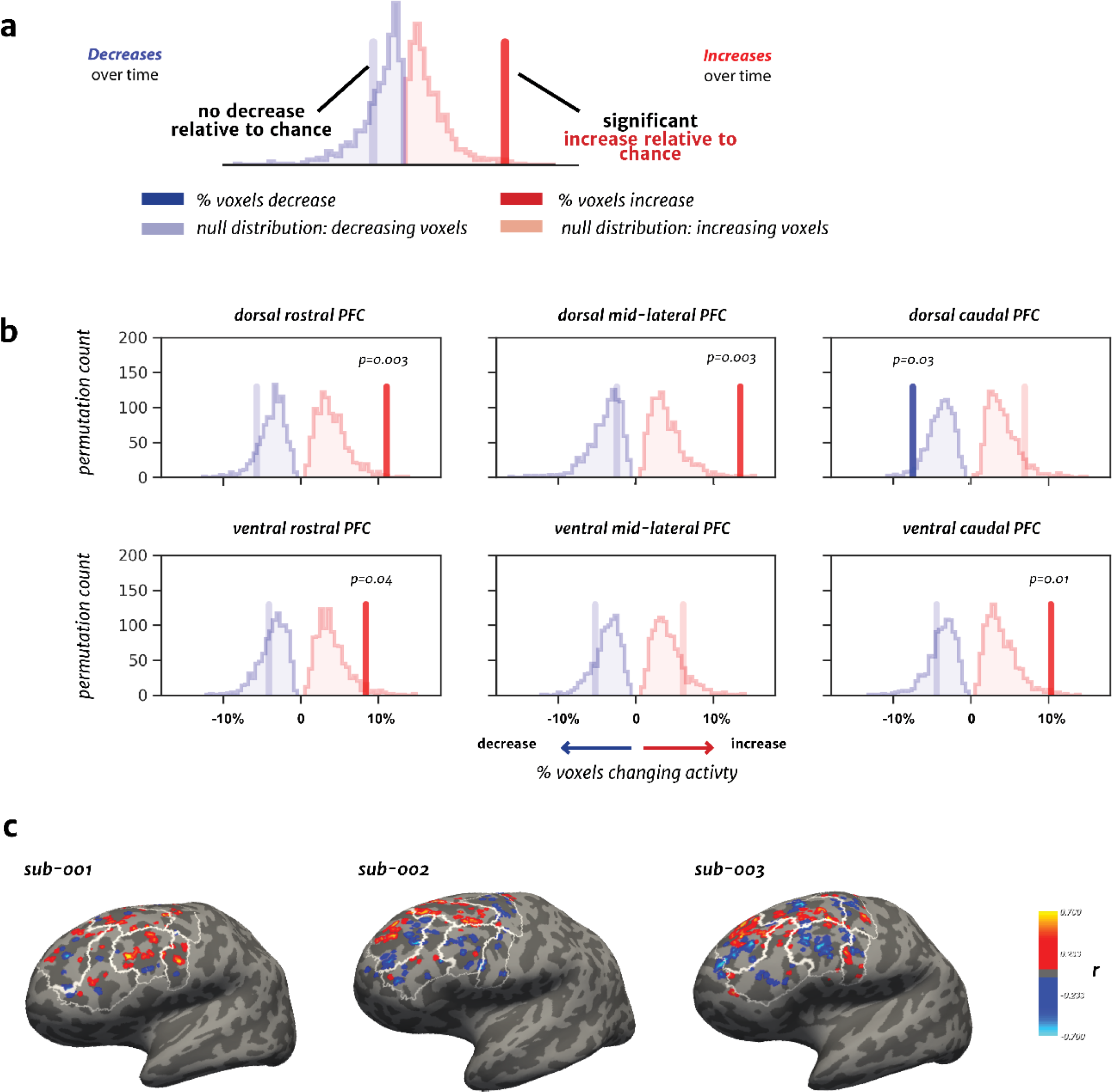
Distribution of WM delay activity patterns in PFC across the course of learning. **(a)** Schematic of distribution of the percentage of voxels with increases (*red; r* > 0) or decreases (*blue; r* < 0) in activity across training (schematic). Significant changes over time are indicated by bolded vertical lines. Null distributions (created by permuting session number in the voxelwise models) are shown in light red and blue. **(b)** The percentage of voxels with significant changes in activity levels across training within each of the six lPFC ROIs. Four ROIs show a significant proportion of voxels with an increase in activity, while only the dorsal caudal PFC also shows a significant proportion of activity decreases. **(c)** Voxelwise maps of activity changes (thresholded at *p* < 0.05) for each participant plotted on their left hemisphere, inflated cortical surface, with the r-value representing the correlation between the predicted and actual activation values shown in Figure 2a. Greater increases are shown with warmer colors, while decreases are shown with cooler colors.

In four lPFC ROIs, a distributed group of voxels increased in WM delay activity with training. That is, a significant percentage of voxels showed increased WM delay activity across the 17 sessions compared to chance (**Figure 3b**; *dorsal rostral: p* = 0.003 [FDR-corrected *p* = 0.018]*, dorsal mid-lateral: p* = 0.003 [FDR-corrected *p* = 0.018]*, ventral rostral: p* = 0.04 [FDR-corrected *p* = 0.11]*, ventral caudal: p* = 0.01 [FDR-corrected *p* = 0.044]; permutation tests). The dorsal mid-lateral and ventral caudal PFC showed the largest percentage of voxels with increasing WM delay activity over months of training (∼25% of voxels). In only the dorsal caudal PFC ROI, a distinct group of voxels exhibited decreased activity (*p* = 0.03), but this did not survive FDR correction ([FDR-corrected *p* = 0.09]). The topography of activation changes over time and mean activity patterns in each participant are shown in **Figure 3** and **SI Figure 5**, and WM delay activity maps for each session are available on NeuroVault (https://neurovault.org/collections/12687/). These observed changes across all of lPFC were specific to the WM delay period, as the encoding (sample) period instead showed widespread decreases in activity with training across all ROIs (**SI Figure 3**).

In summary, repeated task and stimulus exposure was most commonly associated with increased WM delay period activity in a distributed group of voxels across lPFC, suggesting that these areas are more involved in WM maintenance over training. However, this increased activity may stem from the development of selectivity for individual stimuli over time, or a non-specific WM maintenance process that conveys no item-level information content. Therefore, we next tested whether patterns of WM delay activity show evidence for representations of individual trained stimuli and stimulus categories.

### Representational similarity patterns emerge for individual items, stimulus category, and sequence category in WM delay activity

We next tested whether the multivariate activity patterns across populations of voxels develop stimulus specificity over time. We employed a pattern similarity analysis framework (**STAR Methods**) to test whether specific WM representations appear in multi-voxel patterns of delay activity across the course of training. These analyses were designed to test directed hypotheses about training-related changes in item- and category-level representations. We therefore contrasted specific stimulus-pairs with each other to capture various levels of representation: individual items, training category (trained vs. novel items), and sequence membership category. We estimated the similarity of representations across individual stimuli (matrices shown in **Figure 4**; **STAR Methods**) by computing correlations between the WM delay period activity patterns for each stimulus. We then created several models to capture hypothesized levels of representational information (*item-level, category-level, sequence category*) and tested how well the observed similarity patterns matched the idealized models, producing a measure of representational “pattern strength” for each ROI in each session (**STAR Methods**; **Figure 4a**). We then tested whether this pattern strength metric for each model changed across sessions using the same logistic modeling (with cross-validation) as in prior analyses. To determine if any pattern similarity effects were specific to the lPFC or would also be reflected in sensory areas, we examined patterns from early visual cortex (V1-V4) and the lateral occipital complex (LOC), a higher-order visual region (**STAR Methods**).

**Figure 4.**
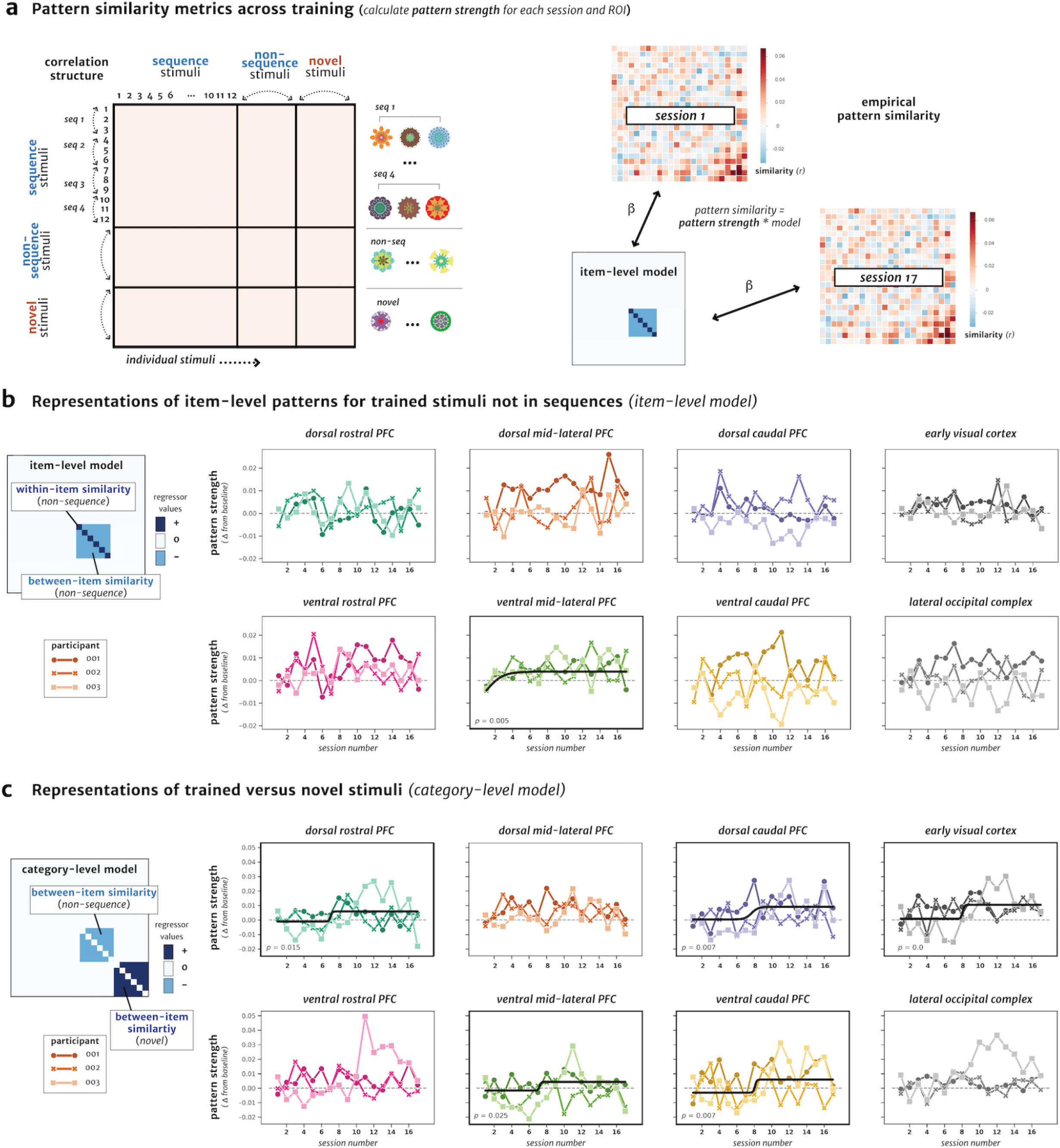
Changes in representational similarity patterns for trained items in WM delay activity. **(a)** Left: Schematic of a WM delay activity pattern similarity matrix across different stimuli. Right: Calculation of the pattern strength metric for each ROI and session by regressing a pattern model against the empirical pattern similarity data. **(b)** Left: Schematic of pattern similarity framework for the *item-level* model, where an interaction between on- (dark blue, positive values) versus off-diagonal (light blue, negative values) correlations among non-sequence trained stimuli serves as a measure of item-level representation. Right: Plots of the pattern strength across sessions for each ROI, as assessed by the model fit for the on- versus off-diagonal interaction. For visualization, all ROIs with significant changes in pattern strength across sessions (after FDR correction) are indicated with a *p*-value and bolded plot border. Pattern strength is plotted and fit with logistic models after subtracting baseline values (**STAR Methods**: *Statistical methods*). Each line shading color and dot style represents one of the three individual participants. **(c)** Same as in **(b)**, but instead testing the category-level model for an interaction between trained (light blue, negative values) versus novel (dark blue, positive values) off-diagonal stimulus correlations.

First, we tested whether distinct representations of individual WM items would emerge across training in lPFC or visual ROIs. We operationalized an *item-level* model for individual stimulus representations by testing for greater within-item pattern similarity (maintenance of the same trained stimulus across different trials, on-diagonal values in correlation matrix) compared to between-item similarity (maintenance of different trained stimuli, off-diagonal correlations), as schematized in **Figure 4b** (*left*). In order to provide the most straightforward and interpretable analysis of item-level representations, we focused on trained items that were *not* part of learned sequences. Analyzing these six stimuli (for each participant) avoids the potential confound that items in temporal sequences may restructure and develop more integrated or differentiated representations over time (Sakai & Miyashita, 1991; Schapiro et al., 2012; Schlichting et al., 2015). In this *item-level* model, higher pattern strength values correspond to stronger representations of individual items in WM (and differentiation from the other trained stimuli). This represents a critical test for the prediction of greater item-level selectivity in lPFC with learning.

Pattern strength for the *item-level* model showed a significant increase over time in ventral mid-lateral lPFC (*ventral mid-lateral*: *r* = 0.36, *p* = 0.005 [FDR-corrected *p* = 0.036]; **Figure 4b**, *right*; inflection point = 1.05 sessions) and not in other PFC or visual areas (all FDR-corrected *p-values* > 0.08). That is, patterns of WM delay activity for individual trained items became more robust (reliable across trials) and differentiated from other trained stimuli across learning. To ensure that this result was not driven by the logistic function modeling approach, we confirmed that this increase in pattern strength in ventral mid-lateral PFC was also reliable when using a linear modeling approach, which demonstrated a significant linear increase over sessions (**SI Results**). Finally, to test whether there were reliable differences in the similarity between each condition after learning began to unfold (after the estimated inflection point), we tested the difference between the on-diagonal (within-item) and off-diagonal (between-item) correlation values. This difference was significant in ventral mid-lateral PFC, with individual items becoming more differentiated from other items (*t* = 2.11, *p* = 0.04). These analyses provide evidence for stronger item-specificity or differentiation of item-level representations in lPFC delay activity across the course of training.

We next asked whether WM representations of all items show evidence of neural differentiation over time, or whether this is specific to trained stimuli. If the item-specific representations in lPFC are specific to trained stimuli, then activation patterns between trained stimuli should become less similar (as the items become more identifiable from each other) while those between novel stimuli should not reliably change. We operationalized this comparison with a *category-level* model which tested for an interaction of a decrease in pattern similarity between trained stimuli (that were not part of sequences) relative to the change in similarity between novel stimuli (off-diagonal correlations) as schematized in **Figure 4c** (*left*). In this *category-level* model, higher pattern strength values correspond to a stronger differentiation between trained and novel stimuli across learning. There was a significant increase in pattern strength for the *category-level* model across sessions in multiple lPFC areas (*dorsal rostral*: *r* = 0.30, *p* = 0.015 [FDR-corrected *p* = 0.030]; *dorsal caudal*: *r* = 0.34, *p* = 0.007 [FDR-corrected *p* = 0.019]; *ventral mid-lateral*: *r* = 0.28, *p* = 0.025 [FDR-corrected *p* = 0.039]; *ventral caudal*: *r* = 0.34, *p* = 0.007 [FDR-corrected *p* = 0.019]) and early visual cortex (*early visual*: *r* = 0.47, *p* < 0.001 [FDR-corrected *p* = 0.002]). The range of inflection points was between sessions 7.23-8.14 for these regions (**Figure 4c**, *right*). The category-level model effect also showed reliable differences in the similarity across conditions, with significantly lower between-item similarity for trained stimuli compared to novel stimuli after the inflection point for each region (*dorsal rostral*: *t* = -3.4, *p* = 0.002; *dorsal caudal*: *t* = -4.4, *p* = 0.0001; *ventral mid-lateral*: *t* = -3.2, *p* = 0.003; *ventral caudal*: *t* = -3.7, *p* = 0.001; *early visual*: *t* = -3.9, *p* = 0.0005). We also confirmed that a linear modeling approach showed the increase in *category-level* pattern strength in the dorsal caudal PFC and early visual cortex, but not in the other ROIs (**SI Results**). These pattern similarity analyses reveal different representational information for trained and novel stimuli across learning, such that the distinction between trained (as compared to novel) stimuli becomes increasingly detectable over time.

Finally, we tested whether associations learned in a distinct task context may influence WM maintenance processes, even when they are not task-relevant. In parallel to the WM task, participants learned that a subset of trained stimuli formed high-probability temporal sequences in the SRT task (**SI Figure 1**). Based on classic studies of paired associate learning (Naya et al., 2001; Sakai & Miyashita, 1991) and multivariate representations that are altered by learning (Schapiro et al., 2012; Schlichting et al., 2015), we tested for shared representations across items in the same temporal sequence (higher similarity across items *within* the same sequence vs. *between* sequences). Surprisingly, we found no increases in this representation over time. We did see a decrease in the early visual cortex, such that items in the same sequence became more distinct from each other over time (relative to between-sequence similarity; **SI Figure 4**).

To better understand any potential higher-level representations influenced by sequence learning, we lastly tested whether the organization of stimuli into temporal sequences in the SRT task may have resulted in a shared representations during WM. That is, a representation between stimuli belonging to *any* sequence (regardless of sequence identity, a generalized sequence representation) which is distinct from items that were not part of temporal sequence structure (non-sequence items). This kind of representation would reflect a categorical difference (or “boundary”) between trained items belonging to a sequence versus those without such associations. This coarse-level representation of sequence category structure was operationalized with a *sequence category* model (**Figure 5**, *left*). The model was designed to compare changes in between-item similarity for items in sequences vs. similarity for sequence-to-nonsequence items and potentially reflect a learned sequence category among trained stimuli. Higher pattern strength values in the model correspond to stronger representations for the category of trained items in sequences from the SRT task, compared to the similarity of these items with other trained stimuli that were not part of temporal sequences.

**Figure 5.**
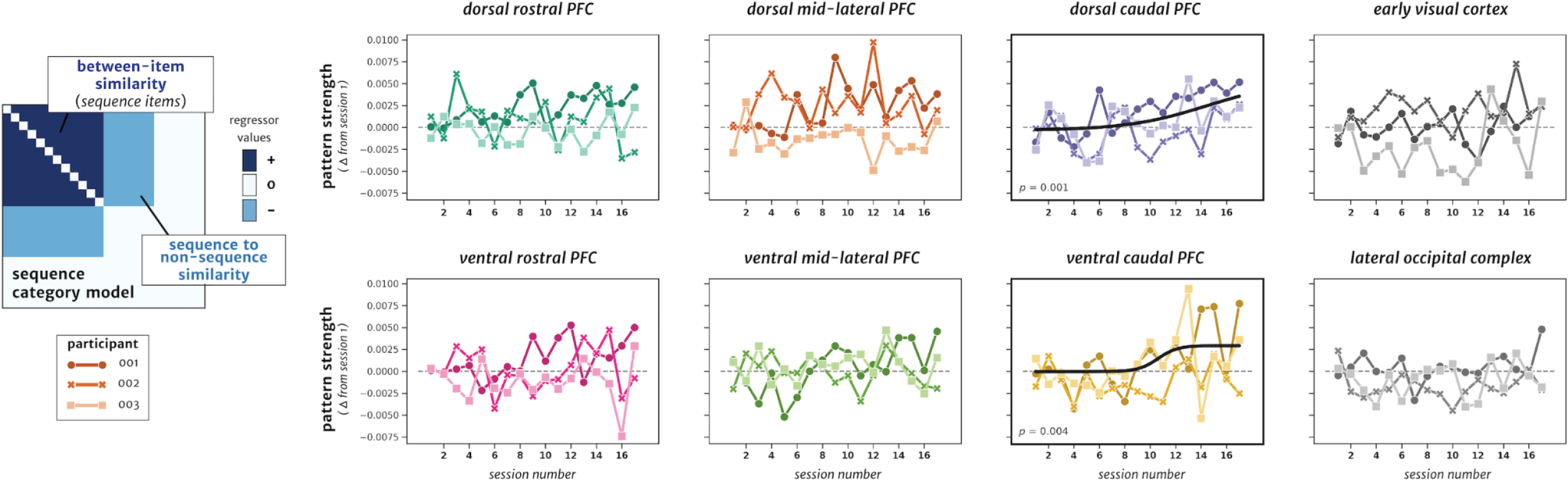
Changes in a categorical sequence representation in WM delay activity. Left: Schematic of the model matrix for pattern similarity between items within trained sequences (dark blue, positive values) compared to trained items not in sequences (light blue, negative values). Right: Plots of the pattern strength for each ROI, at each session, as assessed by the model fit for the *sequence category* model on the left. For visualization, all ROIs with significant changes in pattern strength across sessions (after FDR correction) are indicated with a *p*-value and bolded plot border. Pattern strength is plotted and fit with logistic models after subtracting baseline values (**STAR Methods**: *Statistical methods*). Each line represents one of the three individual participants.

Pattern strength for this *sequence category* model showed a significant increase across sessions in caudal lPFC regions (*dorsal caudal*: *r* = 0.41, *p* = 0.001 [FDR-corrected *p* = 0.011]; *ventral caudal*: *r* = 0.36, *p* = 0.004 [FDR-corrected *p* = 0.017]; **Figure 5**, *right*). The inflection points here were later than any other model (*dorsal caudal*: session 15.2, *ventral caudal*: session 10.5). As in the previous analyses, we also confirmed that the increase in pattern strength for the *sequence category* model in the caudal PFC ROIs was robust when using a different, linear modeling approach (**SI Results**). Finally, there was a reliable mean difference in correlations in the conditions of the sequence category model after learning took place. That is, after the inflection point of the sequence category models, similarity between sequence stimuli was significantly higher than the mean similarity between sequence and non-sequence stimuli (*dorsal cauda*l: *t* = 4.28, *p* = 0.009; *ventral caudal*: *t* = 2.93, *p* = 0.008). Therefore, the sequence category representations were likely driven by higher similarity between trained stimuli that were associated with sequences. Across these analyses that consider associations in the SRT task, stimuli across any learned sequence became more similar to each other over training, relative to stimuli not in sequences, specifically in caudal lPFC regions. These pattern similarity results suggest that learned associations from LTM are reflected in WM delay activity, even when those associations are irrelevant to the WM task goals.

## Discussion

Here, we examined how long-term learning influences lPFC neural representations for WM. Over three months, we extensively trained three human participants on a WM task and a sequence learning (SRT) task, which both employed a unique set of complex, fractal stimuli. We sampled fMRI activity and behavioral performance repeatedly across learning, and found that the distribution and selectivity of lPFC WM delay activity changed with training: more cortical territory was recruited during the WM delay period with learning, and these activity changes coincided with increases in stimulus representations in multivariate patterns (**Figure 6**). Associations between stimuli learned in another task context, although task-irrelevant for WM, also shaped neural representations in lPFC during WM maintenance. In sum, long-term learning changed the distribution and representational structure of lPFC WM delay activity, indicating that the neural mechanisms for WM are influenced by prior experience.

**Figure 6.**
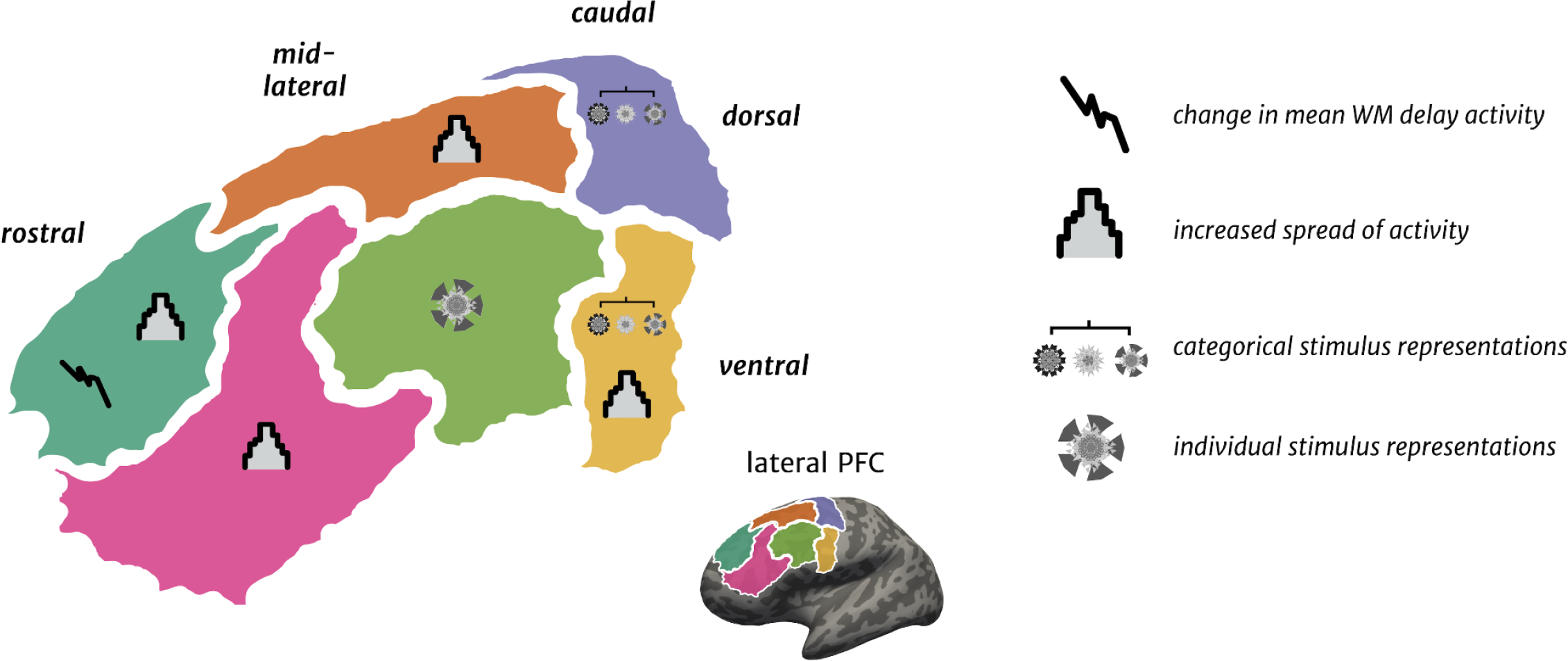
Summary of results. Left: Each lPFC region, with icons depicting which WM delay activity metrics showed training-related changes. Right: Legend for the symbols depicting significant changes in mean WM delay activity magnitude, activity spread within regions, or multivariate representations (from pattern similarity analyses) for sequences and items in WM.

### lPFC - representations or processes?

Early NHP electrophysiological recordings from lPFC revealed neurons that respond to all phases of WM tasks: cue, delay, and response periods (Funahashi et al., 1990). Since then, neurons in NHP lPFC have been shown to encode both stimulus representations (Funahashi et al., 1989; Murray et al., 2017) and cognitive processes, including motor responses, rule learning, and executive control signals (Rigotti et al., 2013; Vallentin et al., 2012; Wallis & Miller, 2003). In contrast, human lPFC shows a relative absence of stimulus specific representations during WM (D’Esposito & Postle, 2015; Harrison & Tong, 2009; Leavitt et al., 2017; Serences, 2016) and human neuroimaging and lesion studies consistently point to lPFC mainly as a source of cognitive control signals (Chatham et al., 2014; Gazzaley & Nobre, 2012; Szczepanski & Knight, 2014). Thus, the role of lPFC function during WM has been unclear across studies. However, NHP and human studies are characterized by stark differences in training regimes before neural recordings take place (Berger et al., 2018; Birman & Gardner, 2016; Sarma et al., 2016). Therefore, we reasoned that differences in task and stimulus experience may underlie the discrepant conclusions about lPFC function.

To directly test the influence of training on WM and lPFC function, we scanned participants across three months of repeated WM task and stimulus exposure. As training progressed, we found that stimulus specific information was increasingly represented in human lPFC delay activity, akin to patterns more commonly found in NHP studies. Across human studies, WM content is typically more difficult to detect in lPFC relative to visual areas (Bhandari et al., 2018; D’Esposito & Postle, 2015; Serences, 2016). However, a few prior studies also detect stimulus representations in human lPFC, for instance, for visual orientation stimuli in retinotopically organized areas (Christophel et al., 2012; Ester et al., 2015). Others decode object category information, but not fine-grained feature information, from PFC (Lee et al., 2013). Therefore, it may be that orientation stimuli or object categories are supported by a distributed, large-scale cortical organization that enables some information to be decoded at the coarse level of BOLD fMRI, even without extensive training. Here, we show that individual representations for visually similar, complex images become increasingly detectable in human lPFC, but not visual areas, through long-term learning. One possible explanation for this training-related change is that the WM information only became represented in lPFC with learning. Another possibility is that there were pre-existing WM representations, below the level of fMRI resolution, but learning altered the representational structure into a more detectable state (i.e. present in large-scale patterns). Likewise, in areas that show no detectable item-level or category representations here, WM representations may exist at finer levels of measurement granularity (either at the sub-voxel resolution, or in finer grained patterns that are not present across entire ROIs), or they may have been too noisy to yield reliable trends over time. Thus, null findings for particular regions (visual cortex) or models should be interpreted with caution. Nonetheless, our results suggest that the debate over the role of lPFC in WM may hinge on training. That is, delay period signals reflecting general WM maintenance (*processes*) are present in fMRI activity without extensive training, while responses to individual stimuli (*representations*) emerge in lPFC after long-term learning.

### Implications for models of functional organization of lPFC

The lPFC is organized in a macroscale gradient along the rostral-caudal axis, both functionally (Badre & Nee, 2018; Koechlin et al., 2003) and anatomically (Goulas et al., 2014; J. A. Miller et al., 2021; Wagstyl et al., 2020). More abstract representations are generally encoded more rostrally along the lPFC (Badre & D’Esposito, 2009), with middle frontal areas posited to sit “atop” the hierarchy and provide top-down control signals during complex cognitive tasks (Badre & Nee, 2018; Duverne & Koechlin, 2017; Ito et al., 2017). Here, our data also support a rostral-caudal organization of WM along lPFC: after training, stimulus-specific representational information emerged in only mid-lateral lPFC and categorical representational information in only caudal lPFC areas (**Figure 6**). The timeline of learning also reflected this abstraction, with progressively later inflection points for more categorical representations. While stimulus categories are often more abstract than individual stimuli – and might therefore be expected to engage more rostral regions – the ‘categorical’ model here may instead capture associations with motor planning processes for item sequences learned in the SRT task. This caudal lPFC sequence-level representation is also consistent with NHP electrophysiology studies finding categorical task and rule representations in anatomically homologous premotor areas (Muhammad et al., 2006; Vallentin et al., 2012; Wallis & Miller, 2003). Altogether, our results suggest that distinct levels of representation for learned stimuli during WM may be scaffolded onto an existing rostro-caudal lPFC functional organization.

Here, we show stimulus-specific activity patterns that are not typically detected in human lPFC, but are more akin to typical observations from NHP studies. However, the exact areas of functional homology are often observed to be anatomically distinct across NHP and human lPFC. For example, lesions of caudal precentral areas in human lPFC cause deficits in spatial WM that mirror the effects in NHPs of damage to more anterior, mid-dorsal lPFC areas (Mackey et al., 2016). Here, we show WM stimulus patterns emerge in micro-anatomically similar areas to where NHP recordings also detect WM stimulus information (e.g., Brodmann’s areas 9/46d, 9/46v; (Petrides, 2005)). Long-term learning drives mid-lateral lPFC regions -- that are most often described as a “controller” of task activity in humans -- to represent stimulus-specific WM information. This suggests the intriguing possibility that the storage location of WM representations and site of top-down control signals can occur in the same area depending on learning and task demands. Future work using longitudinal paradigms in NHP studies might also clarify the importance of training, spatial scale, and species differences on WM maintenance processes (Badre et al., 2015; Milham et al., 2018; Song et al., 2021).

### Plasticity of the PFC

The lPFC is critical for flexible cognition. Multiple theories consider the lPFC to have high plasticity, with activity patterns and representations that change based on task demands (Duncan, 2001; E. K. Miller & Cohen, 2001; Woolgar et al., 2011). However, these patterns of adaptation have not been systematically tracked over time in human lPFC. Some human neuroimaging studies have employed forms of WM training as a route to improve WM and cognition more broadly, but the direction of lPFC change has been inconsistent across studies. Early studies found activation increases in frontal and parietal cortex after WM training (Klingberg, 2010; Olesen et al., 2004), but recent aggregations of WM training studies roughly show activation decreases for studies with shorter training times (∼minutes-hours) and increases for longer training (∼days-weeks) (Buschkuehl et al., 2012, 2014). These studies only sparsely sample neuroimaging data, and have thus been unable to track learning across time, or to examine the effects of stimulus experience and context on the neural mechanisms of WM maintenance. Here we densely sampled neuroimaging and behavioral data across training, and we show progressive increases in both lPFC activity as well as multiple levels of stimulus representational information.

Recently, NHP electrophysiology studies have observed changes after training in the selectivity and magnitude of both single-unit and population spiking during WM (Dang et al., 2021; Meyer et al., 2011; Qi et al., 2019; Tang et al., 2022). Long-term representations of learned stimulus categories are also detectable in frontal and temporal cortex using NHP fMRI (Ghazizadeh et al., 2018), which may serve as a bridge between single-unit and human neuroimaging data. Here, we attempted to approximate the task difficulty and timeline of learning in NHP studies, and we chose a stimulus set of complex, novel fractals with which participants would have no prior experience. The present data shows changes in human lPFC with extensive learning that may parallel changes in activity patterns and representations observed in NHP electrophysiology studies after WM training (Qi et al., 2019; Riley et al., 2018). However, it is difficult to fully bridge across discrepancies in measurement techniques and species: While BOLD fMRI signals can correlate with representations detected via multi-unit activity and high-gamma LFP (Klink et al., 2021; Manea et al., 2022), there is a complicated relationship between spiking activity, LFP signals, and BOLD measurements (Mukamel et al., 2005; Nir et al., 2007; Shi et al., 2017). Ultimately, paradigms using identical tasks, training timelines, and stimuli will be needed to compare the effects of learning on WM neural data between NHP and humans.

The neurophysiological mechanisms underlying such changes in activity patterns with training in both the present data and previous studies are difficult to ascertain. For example, changes in dopaminergic signaling and receptor sensitivity, along with correlated firing across neurons, may all play a mechanistic role in the activation and selectivity increases of lPFC neuronal populations with training (Constantinidis & Klingberg, 2016; Riley et al., 2018; Vijayraghavan et al., 2007). These previous effects have been greatest in mid- and anterior dorsal areas of lPFC, mirroring the organization of emerging stimulus-selective activity patterns that we observed here in mid-lateral lPFC. This lPFC plasticity likely arises from several factors that give the region a high propensity for flexible representations: long-range anatomical connections (Chaudhuri et al., 2015; Y. Wang et al., 2021), status as a hub between cortical networks (Bertolero et al., 2018; Fornito et al., 2019), and a late anatomical development (Garcia et al., 2018; Garcia-Cabezas et al., 2019). Complementing the literature on flexible lPFC activity patterns based on task demands, here we show lPFC plasticity from experience and learning across months.

### Influence of LTM on WM

According to foundational theories, WM and LTM are thought to rely on both different brain areas and neuronal mechanisms for memory storage (Squire & Zola-Morgan, 1991; Warrington & Shallice, 1969; Wickelgren, 1969). Thus, the neural circuitry supporting WM has most often been studied without considering longer term learning and memory effects. However, when WM behavior has been considered in relation to stimulus experience, better WM is observed for familiar, complex stimuli such as Pokémon (Xie & Zhang, 2017), meaningful human faces (Asp et al., 2021; Jackson & Raymond, 2008), and trained geometric shapes (Blalock, 2015). Our findings suggest that these experience-dependent WM behavioral effects are underpinned by malleability of the cortical representations that support WM across learning.

In addition to item-specific patterns, we also found shared WM representations developing for stimuli that were part of temporal sequences in the SRT task, consistent with a “categorical” representation grouping items based on their properties within learned knowledge structures. Long-term memory consolidation is thought to promote the extraction of common features across experiences (McClelland et al., 1995; Winocur & Moscovitch, 2011); thus it is likely that the shared categorical structure emerged as a function of memory consolidation, facilitated by repeated exposure to sequences over time (Antony et al., 2017). The later inflection points of the sequence category models (**Figure 5**) compared to the item-level model (**Figure 4b**) are consistent with a gradual learning and consolidation process facilitating extraction of sequence information over time, which then influenced WM activity patterns. This learning process may have created a semantic-like code for stimuli occupying the same class of patterns over time (sequence stimuli) versus a distinct class of non-sequence stimuli (Binder & Desai, 2011; Eichenbaum, 2017; Nadel & Moscovitch, 1997; Sommer, 2017; Winocur & Moscovitch, 2011). Item-level vs. categorical representations also emerged in different areas of lPFC, suggesting that the activity changes induced by long-term learning obeyed functional axes of lPFC organization (**Figure 6**). For example, sequence category effects in caudal PFC (including premotor areas) may have captured motor planning processes for item sequences. Altogether, the results indicate not only that LTM can share representational formats with WM (Beukers et al., 2021; Lewis-Peacock & Norman, 2014; Nee & Jonides, 2011; Oberauer, 2009), but that long-term learning *changes* how information is represented in WM, even when learned associations are not behaviorally relevant for WM.

### Longitudinal study design interpretations and caveats

In this study, we gained insight into how WM representations change with learning by using a longitudinal, dense sampling design within a targeted group. For example, we show that both item-level and category-level representations become more detectable in human PFC activity patterns with learning, and at different time points in learning. However, the limited sampling of few individuals likely results in findings that do not necessarily generalize to the population of healthy adults. Instead, we offer a detailed characterization of learning effects over months within a specific sample. Therefore, instead of examining variance between participants, we combined the data for participants using a fixed-effects approach to make the most reliable and strongest inference on any effects present in the current sample (rather than the general population, see (Fries & Maris, 2021; Vezoli et al., 2021)). Similar study designs involving a small group of individuals (“deep imaging”) have also been used to show changes in human brain activity and functional organization (Gordon et al., 2017; Naselaris et al., 2021; Newbold et al., 2020). These play a key role in modern neuroscience, placing emphasis on high data quality and within-participant power instead of sampling more individuals to achieve similar levels of statistical power (Gordon et al., 2017; Gratton et al., 2022; Gratton & Braga, 2021; Kragel et al., 2021; Naselaris et al., 2021; Newbold et al., 2020; Popham et al., 2021). The present work leverages this “deep imaging” approach to tackle discrepancies between WM studies.

### Future considerations

Here we show that human lPFC activity patterns gradually change over long-term learning, suggesting that the role of lPFC in WM may shift as stimuli become well-learned and embedded in associative structures. These findings highlight important considerations for conducting and interpreting investigations into WM function. If the neural circuitry for WM is shaped by prior experience, drastically different conclusions can be reached depending on when brain recordings take place relative to training. The timeline of learning is especially important to consider because neuronal ensembles in lPFC demonstrate a remarkable flexibility in activity patterns (e.g., magnitude, timing, and dimensionality) based on behavioral demands across different tasks (Dang et al., 2021; E. K. Miller & Cohen, 2001; E. K. Miller & Fusi, 2013; Stokes et al., 2013; Wasmuht et al., 2018). By implementing a protracted training and recording regime in humans, our data show that long-term learning sculpts neural representations during WM. These data offer a potential bridge between seemingly incompatible accounts from NHP electrophysiology and human fMRI studies. Moving forward, an accurate understanding of PFC and WM functioning should consider training effects, species differences, and how LTM may be involved.

## Author Contributions

J.A.M., A.K., and A.T. developed the study concept and design with input from M.D.. J.A.M., A.K., and A.T. collected and analyzed the data. J.A.M., A.K., and A.T. drafted the manuscript, and M.D. provided feedback and revisions. All authors approved the final version of the manuscript for submission.

## Acknowledgments

This work was supported by National Institutes of Health (NIH) grants F32MH106280 to A.T., F32MH111204 to A.K., and RO1 MH63901 to M.D, and a Wu Tsai Institute postdoctoral fellowship for J.A.M.. We also thank Ian Ballard and Regina Lapate for their input on data analysis and visualization, and Dan Lurie for assistance with data collection.

## STAR Methods

### RESOURCE AVAILABILITY

#### Lead contact

Further information and requests for resources and reagents should be directed to and will be fulfilled by the Lead Contact, Jacob Miller.

#### Materials Availability

This study did not generate new unique reagents.

#### Data and Code Availability

All neuroimaging data will be openly available in the Brain Imaging Data Structure format ((Gorgolewski et al., 2016); https://bids.neuroimaging.io/) on the OpenNeuro platform upon publication (openneuro.org; (Markiewicz et al., 2021)). Analysis and processing code to reproduce the present results, along with the stimuli, presentation code, and behavioral data may be found on Open Science Framework (OSF) : https://osf.io/

### EXPERIMENTAL MODEL AND SUBJECT DETAILS

#### Human participants

The three study participants were all healthy, adult volunteers. Because of the large amount of MRI data collected and intensive nature of the behavioral training involved, all participants were members of the research team who completed the study over the same time period. While participants had limited experience performing WM and cognitive tasks before, this experience is much smaller relative to the study timeline, and they had no prior experience with this particular task or stimulus set. One participant was a 34-year-old female (sub-001), one was a 25-year-old male (sub-002), and one was a 37-year-old female (sub-003). The University of California, Berkeley Committee for the Protection of Human Subjects (CPHS) approved the study protocol and no participants reported any contraindications for MRI.

### METHOD DETAILS

#### Study design and stimuli

The study was designed to investigate WM behavior and neural representations across a large amount of training on a specific set of stimuli and tasks. To accomplish this, we assigned each participant a unique set of 18 fractal images as their set of trained stimuli. Each image was an algorithmically-generated fractal consisting of multiple colors, and the 18 images for each participant were balanced according to the primary color group of each image (determined using a k-means clustering algorithm on each fractal image in the *sklearn* Python package: https://scikit-learn.org/). These fractals were chosen because they are visually complex, approximately uniform in size, cannot be easily verbalized, have no pre-existing meaning, and similar stimuli have been used in NHP electrophysiology studies of the neural basis of learning (Ghazizadeh et al., 2018; Kim et al., 2015; Sakai & Miyashita, 1991). Because the study participants were also on the research team, we avoided participants gaining any foreknowledge of their training set by generating thousands of initial images and randomly selecting each training set from among these images. Thus, each participants’ first exposure to their training set occurred during the first scanning session. The unique 18 stimuli for each participant were then used for all of the following fMRI and behavioral training sessions, with additional novel stimuli randomly selected each session from the broader set of fractals. Of the 18 fractal stimuli in each participant’s training set, 12 were randomly assigned to be part of four sequences in the SRT task, with each sequence consisting of three fractals and an object image. The sequences were learned over time as part of a serial reaction time (SRT) task. Although sequences were not explicitly instructed, all participants had knowledge of the sequence manipulation; thus, reductions in response time in this task likely reflect both explicit and implicit learning. All tasks were programmed using *Psychtoolbox* functions (Brainard, 1997; http://psychtoolbox.org/) in Matlab (https://www.mathworks.com/), and stimuli were presented on a plain white background [RGB = 255,255,255].

#### Longitudinal training

Across the course of 15 weeks, each participant underwent 24-25 total sessions of fMRI scanning. In the present work, we analyze the first 17 of these fMRI sessions (*Phase 1*) for each participant which took place over ∼3 months (13 weeks) of training. In a second study phase (*Phase 2*) of ∼3 additional weeks, more fractal stimuli were added into the training set (**Figure 1c**), but the results from this phase of the experiment are not reported here. Over the first week, four scans were conducted to ensure that the initial exposure to the tasks and stimuli would be highly sampled. fMRI scanning during subsequent weeks occurred at a rate of approximately 1-2x per week (depending on participant and scanner availability).

To facilitate learning, at-home behavioral training was implemented multiple times per week across the course of the study (**Figure 1c**), where participants completed versions of the WM and sequence learning tasks on home laptop testing setups. Most sessions were completed at the same location for each subject, with a small number completed elsewhere (when traveling, for example). The at-home WM task training data can be found on Open Science Framework.

#### Working memory task

Participants completed a three-alternative forced choice delayed recognition task in each scanning and at-home WM training session (**Figure 1a**). Stimuli included the 18 fractals from the participant’s training set, along with 6 novel fractal images, which were randomly selected for each session. On each trial, a single WM sample stimulus (600 x 600 pixels) was presented in the center of a screen for a 0.5 s encoding period. A fixation cross was then presented for a jittered delay period of 4, 8, or 12 s, with the goal of facilitating WM maintenance processes. A probe display then appeared for a response window of 2 s. The probe display comprised three occluded sections of fractal images (⅙ area of each image) at an equal distance from the center of the screen. Each probe image was masked within a gaussian window of FWHM at ∼⅙ the image size. Participants responded via one of three button presses to indicate which probe image segment matched the stimulus from the beginning of the trial. A fourth button option could be used to indicate a guessing response of “I don’t know”. However, this option was rarely chosen (1.6% of total trials), so we could not examine meaningful changes in this response option across sessions. A sample-matching fractal image was always present in the probe display. One of the other probe stimuli was always a novel (untrained) fractal image randomly selected from the same color group as the sample fractal image. The third probe image was either a novel fractal (50% of trials) or a lure from the set of trained fractal images (50% of trials). The masked section of the fractal images was in the same location for each probe image and randomly chosen from nine different areas on each trial, and the probe position was counterbalanced across trials within a block (**Figure 1a**). After each trial, there was a jittered intertrial interval (ITI) sampled from an exponential distribution (mean = 4 s, range = 1 - 9 s).

In the scanning sessions, participants completed four blocks (scanner runs) of 24 trials, with each trained and novel fractal image presented as the WM sample stimulus once per block, in random order. Each delay length occurred in random order and equally often within a block. For the at-home WM training sessions, participants completed two blocks of 24 trials (**Figure 1c**). The in-scanner display was a back-projected 24 in. screen (1024 x 768) for an approximate ∼47 cm viewing distance, while for at-home training sessions participants used laptop screens of sizes 13.3 in. (1440 x 900) [sub-001], 13.3 in. (2560 x 1600) [sub-002], and 12.5 in. (1920 x 1080) [sub-003].

#### Serial reaction time task

In addition to the WM task, participants completed a serial reaction time (SRT) task before the WM task in each scanning session and during at-home training sessions. This task served to repeatedly expose participants to statistical regularities amongst the trained stimuli, in the form of temporal stimulus sequences. During this task, participants made button presses in response to each stimulus. The stimulus set consisted of the same 18 fractal stimuli shown in the WM task as well as six objects (three animals and three tools) for a total of 24 stimuli. The SRT task consisted of two phases: an initial phase in which stimulus-response mappings were learned), followed by a second phase during which stimulus sequences were present.

The first section of SRT task was implemented in the first two sessions of the study (one fMRI session followed by an at-home behavioral session) during which participants were trained to criterion to associate each of the stimuli with one of four button press responses. Participants were first exposed to their stimulus set during their first scanning session. During every block, each of the 24 stimuli were shown once in a randomized order, with no explicit sequence information present (during the first two sessions). Each stimulus was presented on the screen for 2.3 seconds (followed by a blank screen of 0.7 s between stimuli) with four response options shown as black squares below the stimulus (corresponding to the middle finger of the left hand, ring finger of the left hand, ring finger of the right hand, and middle finger of the right hand). During the first two blocks of the first scanning session, the correct response was highlighted (square corresponding to the response was shown in red instead of black) to allow participants to view the correct response and facilitate learning. Thereafter, participants completed 10 more blocks during which the correct response was not shown but feedback was provided (when a correct response was made the square turned blue and incorrect responses were indicated by the selected option turning red with feedback lasting for 200 ms). After the first scanning session, participants performed an at-home session to ensure the learning of stimulus-response mappings. Participants completed a minimum of five blocks of the task, and continued until a criterion of 80% accuracy at the item-level was reached (>=80% of correct first responses for all stimuli across all blocks; 7 - 15 blocks of training were required to reach criterion). The stimulus-response mappings remained constant throughout the study.

After the completion of training to criterion, temporal sequences of stimuli were embedded in the SRT task, beginning in the second fMRI session. Of the 24 trained stimuli (18 fractals and six objects), 16 stimuli were assigned to form four distinct sequences, with each sequence containing three fractals followed by an object (**Figure 1b**). As in the initial section of this task, each stimulus was shown once during each block (set of 24 trials) and the four response options were indicated below the stimulus as four black squares. Participants were instructed to press the appropriate button for each stimulus. Each stimulus was shown for 1.95 s (fMRI sessions) or 1.8 s (behavioral sessions) followed by a blank screen for 400 ms. Sequences were presented in a probabilistic manner, such that three of the four sequences were presented in an intact fashion in each block and each sequence was intact on 75% of blocks in each session (i.e. in 12/16 blocks during fMRI sessions). In each block, the order of the presentation of stimuli was randomized with the exception of the presentation of the three intact sequences. Stimuli from the non-intact sequence (one sequence per block) were presented in a random order with the stipulation that at least two stimuli separated the non-intact sequence stimuli. Feedback was provided throughout the experiment as described above in the training to criterion phase. The fMRI sessions contained 18 blocks of the SRT task and the at-home behavioral sessions consisted of 26 blocks. Stimuli were presented in a randomized order (no sequence information was present) during the first two blocks of each session which served to acclimate participants to the task.

#### Object-selective functional localizer task

Functional localizer scans were collected during two separate fMRI sessions for each participant, which occurred after sessions 1 and 5 for sub-001, sessions 1 and 15 for sub-002, and sessions 5 and 14 for sub-003. Participants performed a one-back task while viewing blocks of animals, tools, objects, faces, scenes, and scrambled images. All images were presented on phase scrambled backgrounds. Each block lasted for 16 s and contained 20 stimuli per block (300 ms stimulus presentation followed by a blank 500 ms inter-stimulus interval). Two stimuli were repeated in each block and participants were instructed to respond to stimulus repetitions via button press. Each scan (three scans per session) contained four blocks of each stimulus class, which were interleaved with five blocks of passive fixation.

#### fMRI acquisition

All neuroimaging data were collected on a 3 Tesla Siemens MRI scanner at the UC Berkeley Henry H. Wheeler Jr. Brain Imaging Center (BIC). Whole-brain Blood Oxygen Level-Dependent (BOLD) fMRI (T_2_*-weighted) scans were acquired with a 32-channel RF head coil using a 2x accelerated multiband echo-planar imaging (EPI) sequence [repetition time (TR) = 2 s, echo time = 30.2 ms, flip angle (FA) = 80°, 2.5 mm isotropic voxels, 52 slices, matrix size = 84 x 84]. Anatomical MRI scans were collected at two timepoints across the study and registered and averaged together before further preprocessing. Each T_1_-weighted anatomical MRI was collected with a 32-channel head coil using an MPRAGE gradient-echo sequence [repetition time (TR) = 2.3 s, echo time = 3 ms, 1 mm isotropic voxels]. For each scan, participants wore custom-fitted headcases (caseforge.com) to facilitate a consistent imaging slice prescription across sessions and to minimize head motion during data acquisition (Power et al., 2019).

In each 2-hr scanning session, participants completed the following BOLD fMRI scans: (1) 9 min eyes-closed rest run, (2) three 9 min runs of a 1-back stimulus localizer, (3) three 6 min runs of the SRT task, (4) 9 min eyes-closed rest block, (5) 9-min stimulus localizer block, (6) four 6 min runs of the WM task. The present work focuses on the WM task. In the stimulus localizer scans, participants viewed trained images in isolation in order to characterize how neural representations change over time (results not reported here).

### QUANTIFICATION AND STATISTICAL ANALYSIS

#### fMRI preprocessing

Preprocessing of the neuroimaging data was performed using fMRIPrep version 1.4.0 (Esteban et al., 2018), a Nipype (Gorgolewski et al., 2017) based tool. Each T1w (T1-weighted) volume was corrected for INU (intensity non-uniformity) using *N4BiasFieldCorrection* v2.1.0 (Tustison et al., 2010) and skull-stripped using *antsBrainExtraction.sh* v2.1.0 (using the OASIS template). Brain tissue segmentation of cerebrospinal fluid (CSF), white-matter (WM) and gray-matter (GM) was performed on the brain-extracted T1w using fast (Zhang et al., 2001) (FSL v5.0.9).

Functional data was slice time corrected using 3dTshift from AFNI v16.2.07 (Cox, 1996) and motion corrected using mcflirt (Jenkinson et al., 2002) (FSL v5.0.9). This was followed by co-registration to the corresponding T1w using boundary-based registration (Greve & Fischl, 2009) with 9 degrees of freedom, using flirt (FSL). Motion correcting transformations and BOLD-to-T1w transformation were concatenated and applied in a single step using *antsApplyTransforms* (ANTs v2.1.0) using Lanczos interpolation. Many internal operations of FMRIPREP use Nilearn (Abraham et al., 2014), principally within the BOLD-processing workflow. For more details of the pipeline see https://fmriprep.readthedocs.io/en/latest/workflows.html. Finally, spatial smoothing was only performed in a 4mm FWHM kernel along the cortical surface (https://github.com/mwaskom/lyman/tree/v2.0.0) for the mean univariate activity analysis (**Figure 2**), while all other analyses used unsmoothed data.

#### Region-of-Interest (ROI) selection

To generate cortical surface reconstructions, the T_1_-weighted anatomical MRIs were processed through the FreeSurfer (https://surfer.nmr.mgh.harvard.edu/) *recon-all* pipeline for gray and white matter segmentation (Dale et al., 1999; Fischl, Sereno, & Dale, 1999). To construct the lPFC ROIs, we sampled a recent multimodal areal parcellation of the human cerebral cortex (Glasser et al., 2016) onto each participant’s native anatomical surface via cortex-based alignment (Fischl, Sereno, Tootell, et al., 1999). We merged these smaller parcels on the surface into six different lPFC ROIs (combined bilaterally), with two splits along the rostral-caudal axis and one split along the dorsal-ventral axis (**Figure 2b**). The caudal lPFC ROIs fall along the precentral sulcus and gyrus, with the most rostral ROIs ending in frontopolar cortex around the anterior ends of the inferior and superior frontal sulci. The split between dorsal-ventral ROIs roughly falls along the posterior middle frontal sulci, analogous microstructurally to the principal sulcus of macaques (J. A. Miller et al., 2021; Petrides, 2019), and the ROIs are bounded dorsally by the superior frontal gyrus and ventrally by the inferior frontal gyrus. This lPFC division into six areas was designed to align with NHP electrophysiology studies recording from multiple frontal cortex regions (Riley et al., 2018).

We also constructed two visual ROIs in order to determine if effects were specific to lPFC or also generalized to lower and higher-order visual areas. An early visual cortex ROI combined visual cortical areas V1-V4 for each participant, defined from aligning a probabilistic visual region atlas (L. Wang et al., 2015) onto each subject’s native cortical surface using cortex-based alignment (**Figure 4a**). A higher-order visual ROI for the lateral occipital complex (LOC) was defined from a separate category localizer scanning session [block-level general linear model (GLM) with a contrast of responses of objects > scrambled objects] (see *Object-selective functional localizer task*). Voxel responses were thresholded at *p* < .0001 and the ROI was restricted to voxels reaching this statistical threshold on the lateral surface of the occipital cortex and the posterior portion of the fusiform gyrus (Schwarzlose et al., 2008) .

#### Mean WM delay activity across training

We constructed a separate event-related GLM in SPM12 (https://www.fil.ion.ucl.ac.uk/spm/) for each participant and session in order to compare activity levels across training. Separate boxcar regressors were constructed for the encoding (0.5 s), delay (4, 8, or 12 s), and probe (2 s) periods of the WM task, and all regressors were convolved with a standard double-gamma hemodynamic response function (HRF). Separate task event regressors were created for trained and novel fractals. For the session-level GLMs, all four WM task runs in each session were concatenated with the *spm_fmri_concatenate* function. Six rigid-body motion parameters were included as nuisance regressors, along with high-pass filtering (HPF) of 128s to capture low-frequency trends as implemented in SPM12 (https://www.fil.ion.ucl.ac.uk/spm/). Voxelwise *beta*-coefficient and *t*-statistic maps were then calculated for WM delay (delay > fixation) periods, selecting regressors for trials across all three delay lengths. We analyzed changes in mean WM delay activity (beta coefficients) over learning with logistic models using mean activity in each ROI as the outcome variable and session number as the predictor (*Statistical methods*). These analyses were performed in two broad groups of voxels: (1) for the mean activity of voxels within the peak activation for each ROI (thresholding the maps for each participant and session at *t* > 2.5) and (2) for the mean activity of all voxels in each ROI (without any thresholding).

For each ROI, we also calculated the time course of activation across the encoding, delay, and probe period for trials with 12 s delay periods (**SI Figure 6**). Activation was plotted at each TR within the trial and normalized relative to a baseline of the mean signal during the ITI period.

#### Voxelwise regression analysis (recruitment of voxels across training)

To ask whether voxels showed changes in activity across training, we performed voxelwise logistic modeling on the *beta-*coefficient values from the above GLMs (*Mean WM delay activity across training*) across sessions (**Figure 2a**, **Figure 3**). Separate voxelwise models were run on WM encoding and delay period activation to characterize changes in each phase of the WM task separately. For each participant and lPFC region, a logistic function was fit to model changes in activation across sessions. Goodness-of-fit of this model was assessed via the correlation between actual and predicted data (using 6-fold cross-validation, see *Statistical methods*). Here, positive values indicate an increase in activity across sessions and negative values indicate a decrease in activity across sessions. After thresholding the voxelwise *r*-value maps (*p* < 0.05, see **Figure 3c** for maps for each participant), we then calculated the proportion of voxels in each ROI showing an increase or decrease in activity across sessions and averaged this value across participants. This generated a measure of how many voxels in an ROI change their activity over time, without requiring overlap of the specific voxels showing changes across participants. To determine if the proportion of voxels showing an increase or decrease in activity across sessions was different than chance (*p* < 0.05 / 2 = 2.5% false-alarm rate for increases or decreases), we constructed permuted null distributions of the proportion of increasing and decreasing voxels in each ROI. In each of 1,000 permutations, session number was randomly shuffled, the same logistic model fitting procedure was performed and regression of predicted activity onto activity across sessions was re-computed, and the proportion of voxels showing increases and decreases in activity (mean across participants) was stored to create null distributions. The true proportion of increasing and decreasing voxels across participants (dark lines in **Figure 3b**) was then compared to the null distributions obtained from permuting session labels to estimate *P*-values representing statistical significance of the proportion of voxels that changed over time.

#### Representational similarity analyses

In order to determine if WM delay activity showed representations for any specific fractal stimuli we obtained single-trial level voxelwise activity maps by constructing separate least-squares-all (LSA) GLMs for each run, session, and participant (Mumford et al., 2012). Here, GLMs were constructed separately for each run in order to estimate pattern similarity between different runs, so that correlation measures aren’t influenced by temporal autocorrelation within each functional scan (Mumford et al., 2014; Zeithamova et al., 2017). In each run-level GLM, the WM delay period events for each of the 24 unique stimuli were modeled as separate boxcar regressors (collapsed across delay lengths) and convolved with a HRF. The combined WM encoding (0.5 s) and probe (2 s) events were included as nuisance regressors, again split by trained and novel stimuli. Six rigid-body motion parameters were also included as nuisance regressors, along with high-pass filtering (HPF) of 128s to capture low-frequency trends. Voxelwise *beta*-coefficient maps from each trial were then used in the pattern similarity analyses.

Before estimating pattern similarity of the delay period activity in each ROI, we applied a multivariate noise decomposition algorithm to the single-trial WM delay period responses (Walther et al., 2016). This process used the time-series of residuals from the LSA GLM for each run to account for noise variance within each ROI, resulting in activation patterns that are less biased by the noise structure. Then, for each session, we calculated between-run correlations (similarity) between the trials for all stimuli (18 trained, 6 novel fractals) across all six run-pair combinations. Correlation values were Fisher-z transformed, and then the mean of the between-run correlations (across run-pairs) generated a representational similarity or correlation matrix (**Figure 5a**). One total run across all sessions and participants was removed from calculation of between-run correlations because of a visual MR artifact (present in the raw functional data). To test for distinct representational structures in WM delay period patterns, we operationalized each of four potential representations as specific predictors of pattern similarity and then analyzed how the strength of each model changed across training. Each representational structure was coded using values of (*1, -1*) for specific stimulus pairs, and negative values were then re-coded such that the regressor values across all conditions summed to zero (i.e. equal weighting was given to positive and negative conditions). After constructing these matrices, they were then used as predictors of the similarity values (Fisher z-transformed pearson correlation), resulting in a model fit (“pattern strength”) for each representational structure. This procedure was performed for each session, participant, and ROI.

First, we constructed an *item-level* model for individual stimulus representations by comparing the on-diagonal correlations (between trials featuring the same stimulus) and off-diagonal correlations for the six trained stimuli not included in any of the learned sequences (**Figure 4b**). Second, we operationalized a category-level model by testing for an interaction in the off-diagonal correlations among all pairs of 18 trained (**Figure 4c**, dark blue) stimuli and the six novel (**Figure 4c**, light blue) stimuli within each session. Finally, we constructed a separate model to test for representations of stimulus sequences from the SRT task. A *sequence category* model tested for an interaction in the similarity of stimuli between *different* sequences (**Figure 5**), compared to a baseline of the correlations between stimuli in sequences to the trained stimuli not in sequences. A final follow-up model directly tested within versus between-sequence stimulus correlations in a sequence identity model, with no differences found across conditions (**SI Figure 4**). For the analysis of off-diagonal correlations among trained stimuli in **Figure 4a**, we excluded the correlations between stimulus pairs within the same sequence from the SRT task.

To determine if there were changes in pattern similarity across training, we used fixed-effects logistic models with the *beta* values from the representational structure matrix regressor (“pattern strength”) as the outcome variable and session number as a predictor. For all models, ROIs with a significant change in the pattern strength across training (significant correlation between the predicted values from the logistic model and the actual data via cross-validation, see *Statistical methods*) are bolded in **Figure 4** and **Figure 5**. We also included early visual and lateral occipital ROIs in the pattern similarity analyses to determine what representational changes are specific to the PFC versus early and higher-order sensory areas.

#### Statistical methods

All changes across training were analyzed using a fixed-effects, logistic model approach, implemented in Scipy’s (https://www.scipy.org/) *optimize.curve_fit* function with four free parameters fit to the data: a slope (*k*), inflection point (*x_T_*), and baseline (*b*) and asymptote (*L*) values (where *x* is session number and *Y* is the outcome variable, such as activation level or pattern strength).

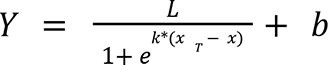

The free parameters were fit within ranges of -10 to 17 for *x_T_*(permitting values before the first session allows a function with relatively small changes in *Y* that occur early in learning and reach a plateau quickly, initial parameter value of median session number), 0 to 10 for *k* (initial value = 0.1), and dynamically adjusted from 2*min(*Y*) to 2*max(*Y*) for *b* (initial value = 0) and *L* (initial value = mean *Y*-value for 20% of latest data points). Code for the logistic fits is available here: https://github.com/arielletambini/logistic-model-fitting

The baseline (mean of session 1 and 2 value) was also subtracted before as part of fitting the model to track changes over time, akin to a linear fixed-effects analysis with a constant regressor for each subject. To avoid overfitting and best capture reliable trends across sessions, we implemented a 6-fold cross-validation scheme during model fitting. Unlike a typical cross-validation procedure in which each fold of the cross-validation uses contiguous data points, here each fold was evenly staggered across the course of learning (session number) in order to not systematically miss one portion of the longitudinal dataset, which could poorly capture learning-related variability. On each fold of cross-validation, either three (5 out of 6 folds) or two (1 of 6 folds) data points were left out of the model fitting procedure, and the predicted values of the held out sessions were stored. Changes over time were assessed by correlating the held out predicted logistic model values from each fold of cross-validation to the actual data values across all sessions (**Figure 2a**). Model parameters were the median value across cross-validation folds. In the few cases where the logistic model failed to converge (**Figure 2c**, *right*: univariate changes in dorsal rostral PFC for all voxels), the model was run again without cross-validation to obtain fit parameters and report the model fit. Data were combined across all participants into one model as fixed-effects in order to assess changes over time, which is recommended for studies with a similar sample size, most often from non-human primate electrophysiology studies (Fries & Maris, 2021; Vezoli et al., 2021).

For each of the RSA analyses of pattern strength across learning (**Figure 4**, **Figure 5**), we also analyzed changes within ROIs that showed a significant logistic model fit with a different modeling approach (**SI Results**). Specifically, we implemented an ordinary-least-squares (OLS) regression with *pattern strength* as the outcome measure and *mean-centered session number* as the predictor, along with the square of session number (2nd order polynomial), as predictor variables. Data was combined across all subjects using a fixed-effects factor (constant for each subject).

For all statistical tests across ROIs, false-discovery rate (FDR) correction (*q* = 0.05) was used to adjust for multiple comparisons across the number of ROIs in each analysis (6 or 8). Neuroimaging files were loaded and operated on using the *Nilearn* package (https://nilearn.github.io/; (Abraham et al., 2014)). For all plots, error bands reflect bootstrapped 68% confidence intervals as implemented in the *Seaborn* package (Waskom, 2021).

## References

Abraham, A., Pedregosa, F., Eickenberg, M., Gervais, P., Mueller, A., Kossaifi, J., Gramfort, A., Thirion, B., & Varoquaux, G. (2014). Machine learning for neuroimaging with scikit-learn. Frontiers in Neuroinformatics, 8, 14.

Antony, J. W., Ferreira, C. S., Norman, K. A., & Wimber, M. (2017). Retrieval as a Fast Route to Memory Consolidation. Trends in Cognitive Sciences. https://doi.org/10.1016/j.tics.2017.05.001

Asp, I. E., Störmer, V. S., & Brady, T. F. (2021). Greater Visual Working Memory Capacity for Visually Matched Stimuli When They Are Perceived as Meaningful. Journal of Cognitive Neuroscience, 33(5), 902–918.

Badre, D., & D’Esposito, M. (2009). Is the rostro-caudal axis of the frontal lobe hierarchical? Nature Reviews. Neuroscience, 10(9), 659–669.

Badre, D., Frank, M. J., & Moore, C. I. (2015). Interactionist Neuroscience. Neuron, 88(5), 855–860.

Badre, D., & Nee, D. E. (2018). Frontal Cortex and the Hierarchical Control of Behavior. Trends in Cognitive Sciences, 22(2), 170–188.

Berger, M., Calapai, A., Stephan, V., Niessing, M., Burchardt, L., Gail, A., & Treue, S. (2018). Standardized automated training of rhesus monkeys for neuroscience research in their housing environment. Journal of Neurophysiology, 119(3), 796–807.

Bertolero, M. A., Yeo, B. T. T., Bassett, D. S., & D’Esposito, M. (2018). A mechanistic model of connector hubs, modularity and cognition. Nature Human Behaviour, 2(10), 765–777.

Beukers, A. O., Buschman, T. J., Cohen, J. D., & Norman, K. A. (2021). Is Activity Silent Working Memory Simply Episodic Memory? Trends in Cognitive Sciences, 25(4), 284–293.

Bhandari, A., Gagne, C., & Badre, D. (2018). Just above Chance: Is It Harder to Decode Information from Prefrontal Cortex Hemodynamic Activity Patterns? Journal of Cognitive Neuroscience, 30(10), 1473–1498.

Binder, J. R., & Desai, R. H. (2011). The neurobiology of semantic memory. Trends in Cognitive Sciences, 15(11), 527–536.

Birman, D., & Gardner, J. L. (2016). Parietal and prefrontal: categorical differences? [Review of Parietal and prefrontal: categorical differences?]. Nature Neuroscience, 19(1), 5–7.

Blalock, L. D. (2015). Stimulus familiarity improves consolidation of visual working memory representations. Attention, Perception & Psychophysics, 77(4), 1143–1158.

Borders, A. A., Ranganath, C., & Yonelinas, A. P. (2021). The hippocampus supports high-precision binding in visual working memory. Hippocampus. https://doi.org/10.1002/hipo.23401

Brady, T. F., Störmer, V. S., & Alvarez, G. A. (2016). Working memory is not fixed-capacity: More active storage capacity for real-world objects than for simple stimuli. Proceedings of the National Academy of Sciences of the United States of America, 113(27), 7459–7464.

Buschkuehl, M., Hernandez-Garcia, L., Jaeggi, S. M., Bernard, J. A., & Jonides, J. (2014). Neural effects of short-term training on working memory. Cognitive, Affective & Behavioral Neuroscience, 14(1), 147–160.

Buschkuehl, M., Jaeggi, S. M., & Jonides, J. (2012). Neuronal effects following working memory training. Developmental Cognitive Neuroscience, 2 Suppl 1, S167–S179.

Chatham, C. H., Frank, M. J., & Badre, D. (2014). Corticostriatal output gating during selection from working memory. Neuron, 81(4), 930–942.

Chaudhuri, R., Knoblauch, K., Gariel, M. A., Kennedy, H., & Wang, X. J. (2015). A Large-Scale Circuit Mechanism for Hierarchical Dynamical Processing in the Primate Cortex. Neuron, 88(2), 419–431.

Christophel, T. B., Hebart, M. N., & Haynes, J. D. (2012). Decoding the contents of visual short-term memory from human visual and parietal cortex. The Journal of Neuroscience: The Official Journal of the Society for Neuroscience, 32(38), 12983–12989.

Christophel, T. B., Klink, P. C., Spitzer, B., Roelfsema, P. R., & Haynes, J. D. (2017). The Distributed Nature of Working Memory. Trends in Cognitive Sciences, 21(2), 111–124.

Constantinidis, C., Funahashi, S., Lee, D., Murray, J. D., Qi, X.-L., Wang, M., & Arnsten, A. F. T. (2018). Persistent Spiking Activity Underlies Working Memory. The Journal of Neuroscience: The Official Journal of the Society for Neuroscience, 38(32), 7020–7028.

Constantinidis, C., & Klingberg, T. (2016). The neuroscience of working memory capacity and training. Nature Reviews. Neuroscience, 17(7), 438–449.

Cox, R. W. (1996). AFNI: software for analysis and visualization of functional magnetic resonance neuroimages. Computers and Biomedical Research, an International Journal, 29(3), 162–173.

Curtis, C. E., & Sprague, T. C. (2021). Persistent Activity during Working Memory from Front to Back. In bioRxiv (p. 2021.04.24.441274). https://doi.org/10.1101/2021.04.24.441274

Dale, A. M., Fischl, B., & Sereno, M. I. (1999). Cortical surface-based analysis. I. Segmentation and surface reconstruction. NeuroImage, 9(2), 179–194.

Dang, W., Jaffe, R. J., Qi, X.-L., & Constantinidis, C. (2021). Emergence of Nonlinear Mixed Selectivity in Prefrontal Cortex after Training. The Journal of Neuroscience: The Official Journal of the Society for Neuroscience, 41(35), 7420–7434.

D’Esposito, M., & Postle, B. R. (2015). The cognitive neuroscience of working memory. Annual Review of Psychology, 66, 115–142.

Duncan, J. (2001). An adaptive coding model of neural function in prefrontal cortex. Nature Reviews. Neuroscience, 2(11), 820–829.

Duverne, S., & Koechlin, E. (2017). Rewards and Cognitive Control in the Human Prefrontal Cortex. Cerebral Cortex, 27(10), 5024–5039.

Eichenbaum, H. (2017). Prefrontal-hippocampal interactions in episodic memory. Nature Reviews. Neuroscience. https://doi.org/10.1038/nrn.2017.74

Eriksson, J., Vogel, E. K., Lansner, A., Bergstrom, F., & Nyberg, L. (2015). Neurocognitive Architecture of Working Memory. Neuron, 88(1), 33–46.

Esteban, O., Markiewicz, C., Blair, R. W., Moodie, C., Isik, A. I., Erramuzpe Aliaga, A., Kent, J., Goncalves, M., DuPre, E., Snyder, M., Oya, H., Ghosh, S., Wright, J., Durnez, J., Poldrack, R., & Gorgolewski, K. J. (2018). FMRIPrep: a robust preprocessing pipeline for functional MRI. bioRxiv. https://doi.org/10.1101/306951

Ester, E. F., Sprague, T. C., & Serences, J. T. (2015). Parietal and Frontal Cortex Encode Stimulus-Specific Mnemonic Representations during Visual Working Memory. Neuron, 87(4), 893–905.

Fischl, B., Sereno, M. I., & Dale, A. M. (1999). Cortical surface-based analysis. II: Inflation, flattening, and a surface-based coordinate system. NeuroImage, 9(2), 195–207.

Fischl, B., Sereno, M. I., Tootell, R. B. H., & Dale, A. M. (1999). High-Resolution Intersubject Averaging and a Coordinate System for the Cortical Surface. Human Brain Mapping, 8, 272–284.

Fornito, A., Arnatkevičiūtė, A., & Fulcher, B. D. (2019). Bridging the Gap between Connectome and Transcriptome. Trends in Cognitive Sciences, 23(1), 34–50.

Fries, P., & Maris, E. (2021). What to do if N is two? In arXiv [stat.ME]. arXiv. http://arxiv.org/abs/2106.14562

Fukuda, K., & Woodman, G. F. (2017). Visual working memory buffers information retrieved from visual long-term memory. Proceedings of the National Academy of Sciences, 201617874.

Funahashi, S., Bruce, C., & Goldman-Rakic, P. S. (1989). Mnemonic encoding of visual space in the monkey’s dorsolateral prefrontal cortex. Journal of Neurophysiology, 61.

Funahashi, S., Bruce, C. J., & Goldman-Rakic, P. S. (1990). Visuospatial coding in primate prefrontal neurons revealed by oculomotor paradigms. Journal of Neurophysiology, 63(4), 814–831.

Fuster, J., & Alexander, G. (1971). Neuron activity related to short-term memory. Science, 173, 652–654.

Garcia-Cabezas, M. A., Zikopoulos, B., & Barbas, H. (2019). The Structural Model: a theory linking connections, plasticity, pathology, development and evolution of the cerebral cortex. Brain Structure & Function. https://doi.org/10.1007/s00429-019-01841-9

Garcia, K. E., Robinson, E. C., Alexopoulos, D., Dierker, D. L., Glasser, M. F., Coalson, T. S., Ortinau, C. M., Rueckert, D., Taber, L. A., Van Essen, D. C., Rogers, C. E., Smyser, C. D., & Bayly, P. V. (2018). Dynamic patterns of cortical expansion during folding of the preterm human brain. Proceedings of the National Academy of Sciences of the United States of America, 115(12), 3156–3161.

Gazzaley, A., & Nobre, A. C. (2012). Top-down modulation: bridging selective attention and working memory. Trends in Cognitive Sciences, 16(2), 129–135.

Ghazizadeh, A., Griggs, W., Leopold, D. A., & Hikosaka, O. (2018). Temporal-prefrontal cortical network for discrimination of valuable objects in long-term memory. Proceedings of the National Academy of Sciences of the United States of America, 115(9), E2135–E2144.

Glasser, M. F., Coalson, T. S., Robinson, E. C., Hacker, C. D., Harwell, J., Yacoub, E., Ugurbil, K., Andersson, J., Beckmann, C. F., Jenkinson, M., Smith, S. M., & Van Essen, D. C. (2016). A multi-modal parcellation of human cerebral cortex. Nature, 536(7615), 171–178.

Goldman-Rakic, P. S. (1995). Cellular Basis of Working Memory. Neuron, 14, 477–485.

Gordon, E. M., Laumann, T. O., Gilmore, A. W., Newbold, D. J., Greene, D. J., Berg, J. J., Ortega, M., Hoyt-Drazen, C., Gratton, C., Sun, H., Hampton, J. M., Coalson, R. S., Nguyen, A. L., McDermott, K. B., Shimony, J. S., Snyder, A. Z., Schlaggar, B. L., Petersen, S. E., Nelson, S. M., & Dosenbach, N. U. F. (2017). Precision Functional Mapping of Individual Human Brains. Neuron. https://doi.org/10.1016/j.neuron.2017.07.011

Gorgolewski, K. J., Auer, T., Calhoun, V. D., Cameron Craddock, R., Das, S., Duff, E. P., Flandin, G., Ghosh, S. S., Glatard, T., Halchenko, Y. O., Handwerker, D. A., Hanke, M., Keator, D., Li, X., Michael, Z., Maumet, C., Nolan Nichols, B., Nichols, T. E., Pellman, J., … Poldrack, R. A. (2016). The brain imaging data structure, a format for organizing and describing outputs of neuroimaging experiments. Scientific Data, 3(1), 1–9.

Gorgolewski, K. J., Esteban, O., Ellis, D. G., Notter, M. P., Ziegler, E., Johnson, H., Hamalainen, C., Yvernault, B., Burns, C., Manhães-Savio, A., Jarecka, D., Markiewicz, C. J., Salo, T., Clark, D., Waskom, M., Wong, J., Modat, M., Dewey, B. E., Clark, M. G., … Ghosh, S. (2017). Nipype: a flexible, lightweight and extensible neuroimaging data processing framework in Python. 0.13.1. https://doi.org/10.5281/zenodo.581704

Goulas, A., Uylings, H. B., & Stiers, P. (2014). Mapping the hierarchical layout of the structural network of the macaque prefrontal cortex. Cerebral Cortex, 24(5), 1178–1194.

Gratton, C., & Braga, R. M. (2021). Editorial overview: Deep imaging of the individual brain: past, practice, and promise. Current Opinion in Behavioral Sciences, 40, iii–vi.

Gratton, C., Nelson, S. M., & Gordon, E. M. (2022). Brain-behavior correlations: Two paths toward reliability [Review of *Brain-behavior correlations: Two paths toward reliability*]. Neuron, 110(9), 1446–1449.

Greve, D. N., & Fischl, B. (2009). Accurate and robust brain image alignment using boundary-based registration. NeuroImage, 48(1), 63–72.

Harrison, S. A., & Tong, F. (2009). Decoding reveals the contents of visual working memory in early visual areas. Nature, 458(7238), 632–635.

Hoskin, A. N., Bornstein, A. M., Norman, K. A., & Cohen, J. D. (2019). Refresh my memory: Episodic memory reinstatements intrude on working memory maintenance. Cognitive, Affective & Behavioral Neuroscience, 19(2), 338–354.

Ito, T., Kulkarni, K. R., Schultz, D. H., Mill, R. D., Chen, R. H., Solomyak, L. I., & Cole, M. W. (2017). Cognitive task information is transferred between brain regions via resting-state network topology. Nature Communications, 8(1), 1027.

Jackson, M. C., & Raymond, J. E. (2008). Familiarity enhances visual working memory for faces. Journal of Experimental Psychology. Human Perception and Performance, 34(3), 556–568.

Jenkinson, M., Bannister, P., Brady, M., & Smith, S. (2002). Improved optimization for the robust and accurate linear registration and motion correction of brain images. NeuroImage, 17(2), 825–841.

Kim H. F., Ghazizadeh, A., & Hikosaka, O. (2015). Dopamine Neurons Encoding Long-Term Memory of Object Value for Habitual Behavior. Cell, 163(5), 1165–1175.

Klingberg, T. (2010). Training and plasticity of working memory. Trends in Cognitive Sciences, 14(7), 317–324.

Klink, P. C., Chen, X., Vanduffel, W., & Roelfsema, P. R. (2021). Population receptive fields in nonhuman primates from whole-brain fMRI and large-scale neurophysiology in visual cortex. eLife, 10. https://doi.org/10.7554/eLife.67304

Koechlin, E., Ody, C., & Kouneiher, F. (2003). The architecture of cognitive control in the human prefrontal cortex. Science, 302(5648), 1181–1185.

Kragel, P. A., Han, X., Kraynak, T. E., Gianaros, P. J., & Wager, T. D. (2021). Functional MRI Can Be Highly Reliable, but It Depends on What You Measure: A Commentary on Elliott et al. (2020) [Review of *Functional MRI Can Be Highly Reliable, but It Depends on What You Measure: A Commentary on Elliott et al. (2020)]*. Psychological Science, 32(4), 622–626.

Lara, A. H., & Wallis, J. D. (2015). The Role of Prefrontal Cortex in Working Memory: A Mini Review. Frontiers in Systems Neuroscience, 9, 173.

LaRocque, J. J., Lewis-Peacock, J. A., & Postle, B. R. (2014). Multiple neural states of representation in short-term memory? It’s a matter of attention. Frontiers in Human Neuroscience, 8.

Leavitt, M. L., Mendoza-Halliday, D., & Martinez-Trujillo, J. C. (2017). Sustained Activity Encoding Working Memories: Not Fully Distributed. Trends in Neurosciences, 40(6), 328–346.

Lee, S. H., Kravitz, D. J., & Baker, C. I. (2013). Goal-dependent dissociation of visual and prefrontal cortices during working memory. Nature Neuroscience, 16(8), 997–999.

Lewis-Peacock, J. A., & Norman, K. A. (2014). Competition between items in working memory leads to forgetting. Nature Communications, 5, 5768.

Lorenc, E. S., & Sreenivasan, K. K. (2021). Reframing the debate: The distributed systems view of working memory. Visual Cognition, 1–9.

Mackey, W. E., Devinsky, O., Doyle, W. K., Meager, M. R., & Curtis, C. E. (2016). Human Dorsolateral Prefrontal Cortex Is Not Necessary for Spatial Working Memory. The Journal of Neuroscience: The Official Journal of the Society for Neuroscience, 36(10), 2847–2856.

Manea, A. M. G., Zilverstand, A., Ugurbil, K., Heilbronner, S. R., & Zimmermann, J. (2022). Intrinsic timescales as an organizational principle of neural processing across the whole rhesus macaque brain. eLife, 11. https://doi.org/10.7554/eLife.75540

Markiewicz, C. J., Gorgolewski, K. J., Feingold, F., Blair, R., Halchenko, Y. O., Miller, E., Hardcastle, N., Wexler, J., Esteban, O., Goncavles, M., Jwa, A., & Poldrack, R. (2021). The OpenNeuro resource for sharing of neuroscience data. eLife, 10. https://doi.org/10.7554/eLife.71774

McClelland, J. L., McNaughton, B. L., & O’Reilly, R. C. (1995). Why there are complementary learning systems in the hippocampus and neocortex: insights from the successes and failures of connectionist models of learning and memory. Psychological Review, 102(3), 419–457.

Mendoza-Halliday, D., Torres, S., & Martinez-Trujillo, J. C. (2014). Sharp emergence of feature-selective sustained activity along the dorsal visual pathway. Nature Neuroscience, 17(9), 1255–1262.

Meyer, T., Qi, X.-L., Stanford, T. R., & Constantinidis, C. (2011). Stimulus selectivity in dorsal and ventral prefrontal cortex after training in working memory tasks. The Journal of Neuroscience: The Official Journal of the Society for Neuroscience, 31(17), 6266–6276.

Milham, M. P., Ai, L., Koo, B., Xu, T., Amiez, C., Balezeau, F., Baxter, M. G., Blezer, E. L. A., Brochier, T., Chen, A., Croxson, P. L., Damatac, C. G., Dehaene, S., Everling, S., Fair, D. A., Fleysher, L., Freiwald, W., Froudist-Walsh, S., Griffiths, T. D., … Schroeder, C. E. (2018). An Open Resource for Non-human Primate Imaging. Neuron, 100(1), 61–74.e2.

Miller, E. K., & Cohen, J. D. (2001). AN INTEGRATIVE THEORY OF PREFRONTAL CORTEX FUNCTION. Annual Review of Neuroscience, 24, 167–202.

Miller, E. K., & Fusi, S. (2013). Limber neurons for a nimble mind. Neuron, 78(2), 211–213.

Miller, E. K., Lundqvist, M., & Bastos, A. M. (2018). Working Memory 2.0. Neuron, 100(2), 463–475.

Miller, J. A., Voorhies, W. I., Lurie, D. J., D’Esposito, M., & Weiner, K. S. (2021). Overlooked Tertiary Sulci Serve as a Meso-Scale Link between Microstructural and Functional Properties of Human Lateral Prefrontal Cortex. The Journal of Neuroscience: The Official Journal of the Society for Neuroscience, 41(10), 2229–2244.

Muhammad, R., Wallis, J. D., & Miller, E. K. (2006). A comparison of abstract rules in the prefrontal cortex, premotor cortex, inferior temporal cortex, and striatum. Journal of Cognitive Neuroscience, 18(6), 974–989.

Mukamel, R., Gelbard, H., Arieli, A., Hasson, U., Fried, I., & Malach, R. (2005). Coupling between neuronal firing, field potentials, and FMRI in human auditory cortex. Science, 309(5736), 951–954.

Mumford, J. A., Davis, T., & Poldrack, R. A. (2014). The impact of study design on pattern estimation for single-trial multivariate pattern analysis. NeuroImage, 103, 130–138.

Mumford, J. A., Turner, B. O., Ashby, F. G., & Poldrack, R. A. (2012). Deconvolving BOLD activation in event-related designs for multivoxel pattern classification analyses. NeuroImage, 59(3), 2636–2643.

Murray, J. D., Bernacchia, A., Roy, N. A., Constantinidis, C., Romo, R., & Wang, X. J. (2017). Stable population coding for working memory coexists with heterogeneous neural dynamics in prefrontal cortex. Proceedings of the National Academy of Sciences of the United States of America, 114(2), 394–399.

Nadel, L., & Moscovitch, M. (1997). Memory consolidation, retrograde amnesia and the hippocampal complex. Current Opinion in Neurobiology, 7(2), 217–227.

Naselaris, T., Allen, E., & Kay, K. (2021). Extensive sampling for complete models of individual brains. Current Opinion in Behavioral Sciences, 40, 45–51.

Naya, Y., Yoshida, M., & Miyashita, Y. (2001). Backward spreading of memory-retrieval signal in the primate temporal cortex. Science, 291(5504), 661–664.

Nee, D. E., & Jonides, J. (2011). Dissociable contributions of prefrontal cortex and the hippocampus to short-term memory: evidence for a 3-state model of memory. NeuroImage, 54(2), 1540–1548.

Newbold, D. J., Laumann, T. O., Hoyt, C. R., Hampton, J. M., Montez, D. F., Raut, R. V., Ortega, M., Mitra, A., Nielsen, A. N., Miller, D. B., Adeyemo, B., Nguyen, A. L., Scheidter, K. M., Tanenbaum, A. B., Van, A. N., Marek, S., Schlaggar, B. L., Carter, A. R., Greene, D. J., … Dosenbach, N. U. F. (2020). Plasticity and Spontaneous Activity Pulses in Disused Human Brain Circuits. Neuron, 107(3), 580–589.e6.

Nir, Y., Fisch, L., Mukamel, R., Gelbard-Sagiv, H., Arieli, A., Fried, I., & Malach, R. (2007). Coupling between neuronal firing rate, gamma LFP, and BOLD fMRI is related to interneuronal correlations. Current Biology: CB, 17(15), 1275–1285.

Oberauer, K. (2009). Design for a Working Memory. In The Psychology of Learning and Motivation (pp. 45–100).

Olesen, P. J., Westerberg, H., & Klingberg, T. (2004). Increased prefrontal and parietal activity after training of working memory. Nature Neuroscience, 7(1), 75–79.

Park, S. H., Russ, B. E., McMahon, D. B. T., Koyano, K. W., Berman, R. A., & Leopold, D. A. (2017). Functional Subpopulations of Neurons in a Macaque Face Patch Revealed by Single-Unit fMRI Mapping. Neuron. https://doi.org/10.1016/j.neuron.2017.07.014

Petrides, M. (2005). Lateral prefrontal cortex: architectonic and functional organization. Philosophical Transactions of the Royal Society of London. Series B, Biological Sciences, 360(1456), 781–795.

Petrides, M. (2019). Atlas of the Morphology of the Human Cerebral Cortex on the Average MNI Brain (1st ed.). Elsevier.

Popham, S. F., Huth, A. G., Bilenko, N. Y., Deniz, F., Gao, J. S., Nunez-Elizalde, A. O., & Gallant, J. L. (2021). Visual and linguistic semantic representations are aligned at the border of human visual cortex. Nature Neuroscience, 24(11), 1628–1636.

Power, J. D., Silver, B. M., Silverman, M. R., Ajodan, E. L., Bos, D. J., & Jones, R. M. (2019). Customized head molds reduce motion during resting state fMRI scans. NeuroImage, 189, 141–149.

Qi, X. L., Riley, M. R., & Constantinidis, C. (2019). Working memory capacity is enhanced by distributed prefrontal activation and invariant temporal dynamics. Proceedings of the. https://www.pnas.org/content/116/14/7095.short

Ranganath, C., & Blumenfeld, R. S. (2005). Doubts about double dissociations between short- and long-term memory. Trends in Cognitive Sciences, 9(8), 374–380.

Ranganath, C., Johnson, M. K., & D’Esposito, M. (2003). Prefrontal activity associated with working memory and episodic long-term memory. Neuropsychologia, 41(3), 378–389.

Riggall, A. C., & Postle, B. R. (2012). The relationship between working memory storage and elevated activity as measured with functional magnetic resonance imaging. The Journal of Neuroscience: The Official Journal of the Society for Neuroscience, 32(38), 12990–12998.

Rigotti, M., Barak, O., Warden, M. R., Wang, X.-J., Daw, N. D., Miller, E. K., & Fusi, S. (2013). The importance of mixed selectivity in complex cognitive tasks. Nature, 497(7451), 585–590.

Riley, M. R., Qi, X. L., Zhou, X., & Constantinidis, C. (2018). Anterior-posterior gradient of plasticity in primate prefrontal cortex. Nature Communications, 9(1), 3790.

Romo, R., Brody, C. D., Hernández, A., & Lemus, L. (1999). Neuronal correlates of parametric working memory in the prefrontal cortex. Nature, 399(6735), 470–473.

Sakai, K., & Miyashita, Y. (1991). Neural organization for the long-term memory of paired associates. Nature, 354(6349), 152–155.

Sarma, A., Masse, N. Y., Wang, X.-J., & Freedman, D. J. (2016). Task-specific versus generalized mnemonic representations in parietal and prefrontal cortices. Nature Neuroscience, 19(1), 143–149.

Schapiro, A. C., Kustner, L. V., & Turk-Browne, N. B. (2012). Shaping of object representations in the human medial temporal lobe based on temporal regularities. Current Biology: CB, 22(17), 1622–1627.

Schlichting, M. L., Mumford, J. A., & Preston, A. R. (2015). Learning-related representational changes reveal dissociable integration and separation signatures in the hippocampus and prefrontal cortex. Nature Communications, 6, 8151.

Schwarzlose, R. F., Swisher, J. D., Dang, S., & Kanwisher, N. (2008). The distribution of category and location information across object-selective regions in human visual cortex. Proceedings of the National Academy of Sciences of the United States of America, 105(11), 4447–4452.

Serences, J. T. (2016). Neural mechanisms of information storage in visual short-term memory. Vision Research, 128, 53–67.

Shi, Z., Wu, R., Yang, P.-F., Wang, F., Wu, T.-L., Mishra, A., Chen, L. M., & Gore, J. C. (2017). High spatial correspondence at a columnar level between activation and resting state fMRI signals and local field potentials. Proceedings of the National Academy of Sciences of the United States of America, 114(20), 201620520.

Sommer, T. (2017). The Emergence of Knowledge and How it Supports the Memory for Novel Related Information. Cerebral Cortex, 27(3), 1906–1921.

Song, X., García-Saldivar, P., Kindred, N., Wang, Y., Merchant, H., Meguerditchian, A., Yang, Y., Stein, E. A., Bradberry, C. W., Ben Hamed, S., Jedema, H. P., & Poirier, C. (2021). Strengths and challenges of longitudinal non-human primate neuroimaging. NeuroImage, 236, 118009.

Squire, L. R., & Zola-Morgan, S. (1991). The medial temporal lobe memory system. Science, 253(5026), 1380–1386.

Sreenivasan, K. K., Curtis, C. E., & D’Esposito, M. (2014). Revisiting the role of persistent neural activity during working memory. Trends in Cognitive Sciences, 18(2), 82–89.

Stokes, M. G., Kusunoki, M., Sigala, N., Nili, H., Gaffan, D., & Duncan, J. (2013). Dynamic coding for cognitive control in prefrontal cortex. Neuron, 78(2), 364–375.

Supèr, H., Spekreijse, H., & Lamme, V. A. (2001). A neural correlate of working memory in the monkey primary visual cortex. Science, 293(5527), 120–124.

Szczepanski, S. M., & Knight, R. T. (2014). Insights into human behavior from lesions to the prefrontal cortex. Neuron, 83(5), 1002–1018.

Tang, H., Riley, M. R., Singh, B., Qi, X.-L., Blake, D. T., & Constantinidis, C. (2022). Prefrontal cortical plasticity during learning of cognitive tasks. Nature Communications, 13(1), 90.

Tustison, N. J., Avants, B. B., Cook, P. A., Zheng, Y., Egan, A., Yushkevich, P. A., & Gee, J. C. (2010). N4ITK: improved N3 bias correction. IEEE Transactions on Medical Imaging, 29(6), 1310–1320.

Vallentin, D., Bongard, S., & Nieder, A. (2012). Numerical rule coding in the prefrontal, premotor, and posterior parietal cortices of macaques. The Journal of Neuroscience: The Official Journal of the Society for Neuroscience, 32(19), 6621–6630.

Vezoli, J., Vinck, M., Bosman, C. A., Bastos, A. M., Lewis, C. M., Kennedy, H., & Fries, P. (2021). Brain rhythms define distinct interaction networks with differential dependence on anatomy. Neuron. https://doi.org/10.1016/j.neuron.2021.09.052

Vijayraghavan, S., Wang, M., Birnbaum, S. G., Williams, G. V., & Arnsten, A. F. T. (2007). Inverted-U dopamine D1 receptor actions on prefrontal neurons engaged in working memory. Nature Neuroscience, 10(3), 376–384.

Wagstyl, K., Larocque, S., Cucurull, G., Lepage, C., Cohen, J. P., Bludau, S., Palomero-Gallagher, N., Lewis, L. B., Funck, T., Spitzer, H., Dickscheid, T., Fletcher, P. C., Romero, A., Zilles, K., Amunts, K., Bengio, Y., & Evans, A. C. (2020). BigBrain 3D atlas of cortical layers: Cortical and laminar thickness gradients diverge in sensory and motor cortices. PLoS Biology, 18(4), e3000678.

Wallis, J. D., & Miller, E. K. (2003). From rule to response: neuronal processes in the premotor and prefrontal cortex. Journal of Neurophysiology, 90(3), 1790–1806.

Walther, A., Nili, H., Ejaz, N., Alink, A., Kriegeskorte, N., & Diedrichsen, J. (2016). Reliability of dissimilarity measures for multi-voxel pattern analysis. NeuroImage, 137, 188–200.

Wang L., Mruczek, R. E., Arcaro, M. J., & Kastner, S. (2015). Probabilistic Maps of Visual Topography in Human Cortex. Cerebral Cortex, 25(10), 3911–3931.

Wang Y., Royer, J., Park, B.-Y., de Wael, R. V., Lariviere, S., Tavakol, S., Rodriguez-Cruces, R., Paquola, C., Hong, S.-J., Margulies, D., Smallwood, J., Valk, S., Evans, A., & Bernhardt, B. C. (2021). Long-range connections mirror and link microarchitectural and cognitive hierarchies in the human brain. In bioRxiv (p. 2021.10.25.465692). https://doi.org/10.1101/2021.10.25.465692

Warrington, E. K., & Shallice, T. (1969). The selective impairment of auditory verbal short-term memory. Brain: A Journal of Neurology, 92(4), 885–896.

Waskom, M. (2021). seaborn: statistical data visualization. Journal of Open Source Software, 6(60), 3021.

Wasmuht, D. F., Spaak, E., Buschman, T. J., Miller, E. K., & Stokes, M. G. (2018). Intrinsic neuronal dynamics predict distinct functional roles during working memory. Nature Communications, 9(1), 3499.

Wickelgren, W. A. (1969). Sparing of short-term memory in an amnesic patient: implications for strength theory of memory. In Neurocase (Vol. 2, Issue 4, p. 259as – 298). https://doi.org/10.1093/neucas/2.4.259-as

Winocur, G., & Moscovitch, M. (2011). Memory transformation and systems consolidation. Journal of the International Neuropsychological Society: JINS, 17(5), 766–780.

Woolgar, A., Hampshire, A., Thompson, R., & Duncan, J. (2011). Adaptive coding of task-relevant information in human frontoparietal cortex. The Journal of Neuroscience: The Official Journal of the Society for Neuroscience, 31(41), 14592–14599.

Xie, W., & Zhang, W. (2017). Familiarity increases the number of remembered Pokémon in visual short-term memory. Memory & Cognition, 45(4), 677–689.

Yonelinas, A. P. (2013). The hippocampus supports high-resolution binding in the service of perception, working memory and long-term memory. Behavioural Brain Research, 254, 34–44.

Zeithamova, D., de Araujo Sanchez, M. A., & Adke, A. (2017). Trial timing and pattern-information analyses of fMRI data. NeuroImage, 153, 221–231.

Zhang, Y., Brady, M., & Smith, S. (2001). Segmentation of brain MR images through a hidden Markov random field model and the expectation-maximization algorithm. IEEE Transactions on Medical Imaging, 20(1), 45–57.

