## Supplemental Information for "Long-term learning transforms prefrontal cortex representations during working memory"

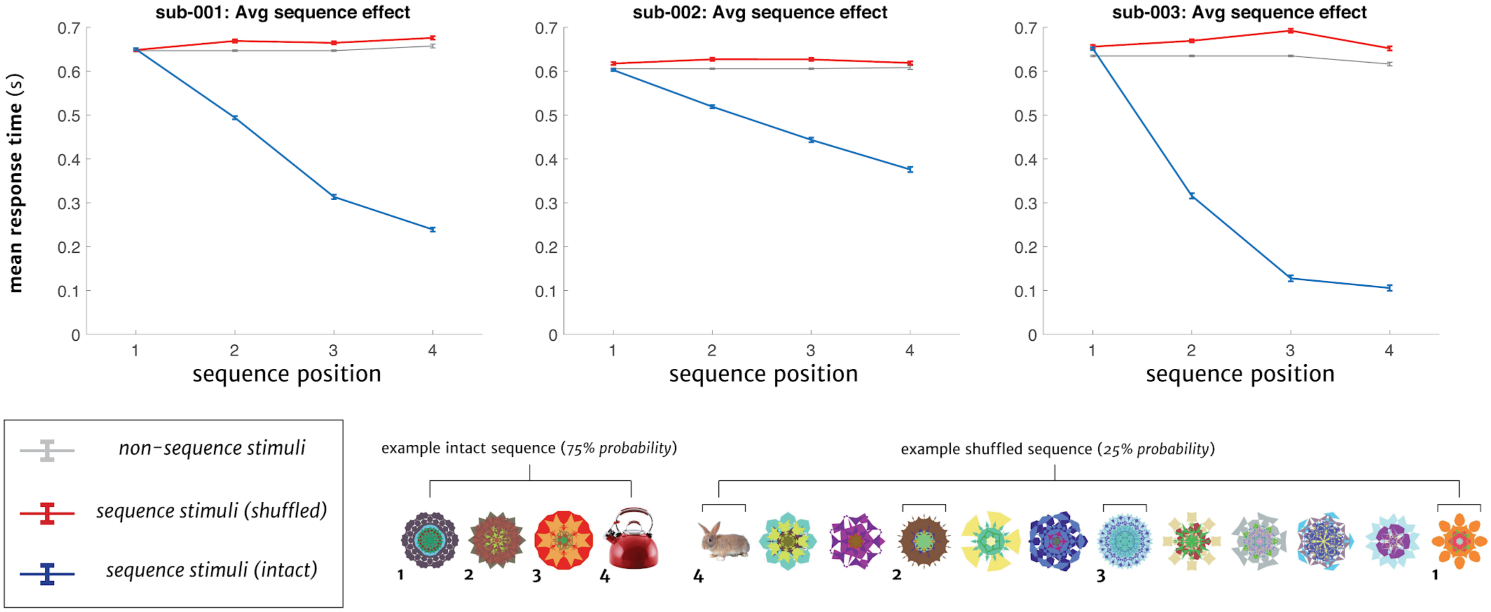


**SI Figure 1. Sequence learning in the Serial Reaction Time (SRT) task.**
Top: Mean response time (s) for correct trials is plotted for each participant across sequence position (1-4) for intact sequences (*blue*), compared to when the same stimuli were shown out of order (*shuffled, red*), and relative to non-sequence stimuli for reference (*gray*). All three participants showed significantly speeded responses across stimuli in intact sequences during fMRI sessions (sub-001, Position 2: $t(15) = -5.50$, $p = 6.1 \times 10^{-5}$; Position 3: $t(15) = -6.29$, $p = 1.4 \times 10^{-5}$; Position 4: $t(15) = -8.58$, $p = 3.6 \times 10^{-7;}$ ; sub-002, Position 2: $t(15) = -7.90$, $p = 1.0 \times 10^{-6}$; Position 3: $t(15) = -7.5$, $p = 1.8 \times 10^{-6}$; Position 4: $t(15) = -9.4$, $p = 1.1 \times 10^{-7}$; sub-003, Position 2: $t(15) = -7.80$, $p = 1.2 \times 10^{-6}$; Position 3: $t(15) = -8.6$, $p = 3.2 \times 10^{-7}$; Position 4: $t(15) = -8.6$, $p = 3.3 \times 10^{-7}$). Bottom: Examples of an intact, left, or shuffled, right, sequence in the SRT task. Intact sequences occurred with higher probability (75%) than shuffled sequences (25%). Error bars represent 68% CI (S.E.M.).

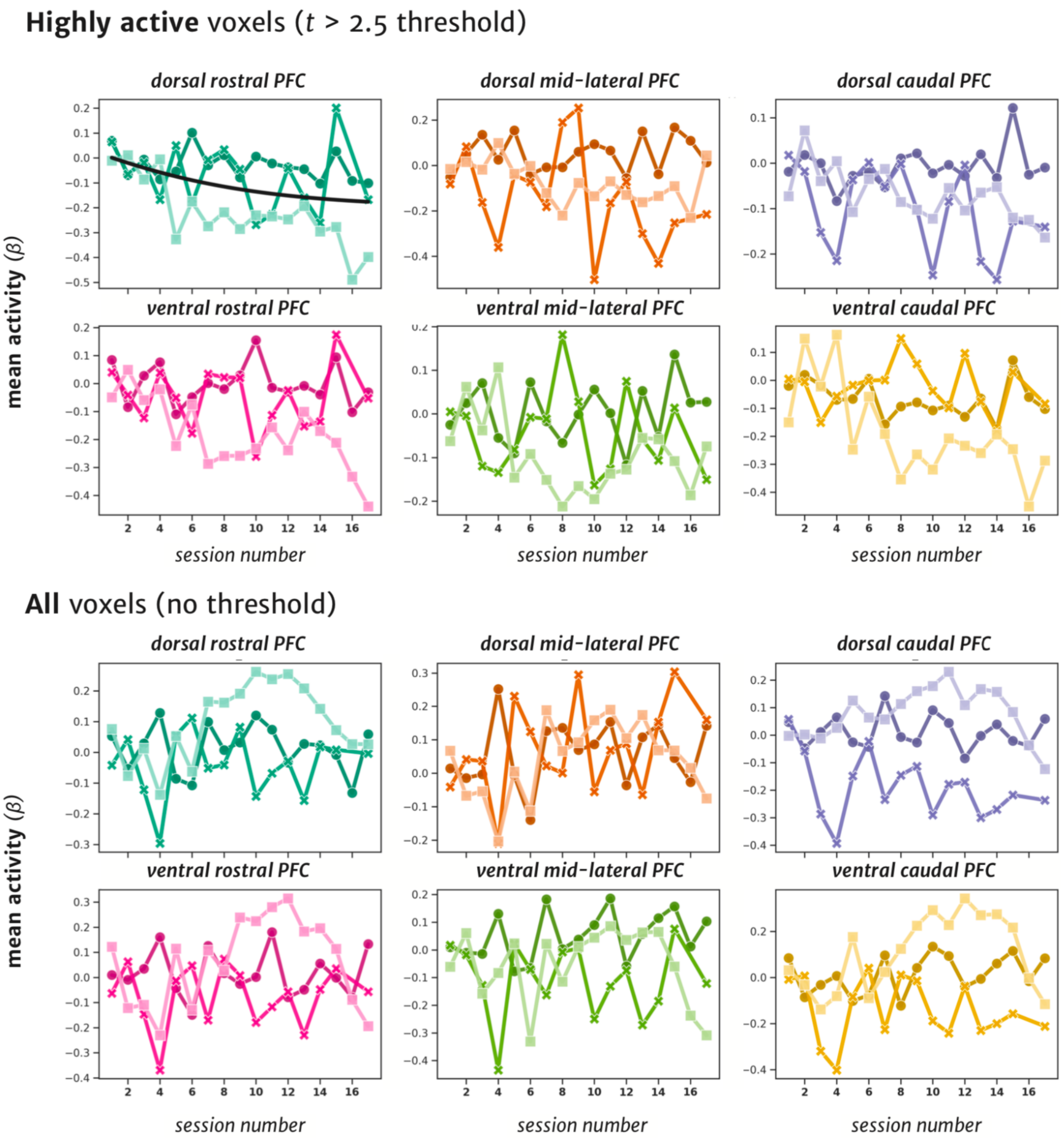


**SI Figure 2. Mean WM delay activity in all lPFC regions across training.**
Top: Mean activity for each fMRI session during the WM delay period for highly active voxels in each lPFC ROI, thresholded at $t > 2.5$ (baseline subtracted). Specific to dorsal rostral PFC (*green*), there was a mean decrease in WM delay activity in the ROI across sessions. Bottom: For all voxels (unthresholded) in an ROI, there were no increases in WM delay across any PFC areas. Bold lines represent logistic model curves from significant model fits.

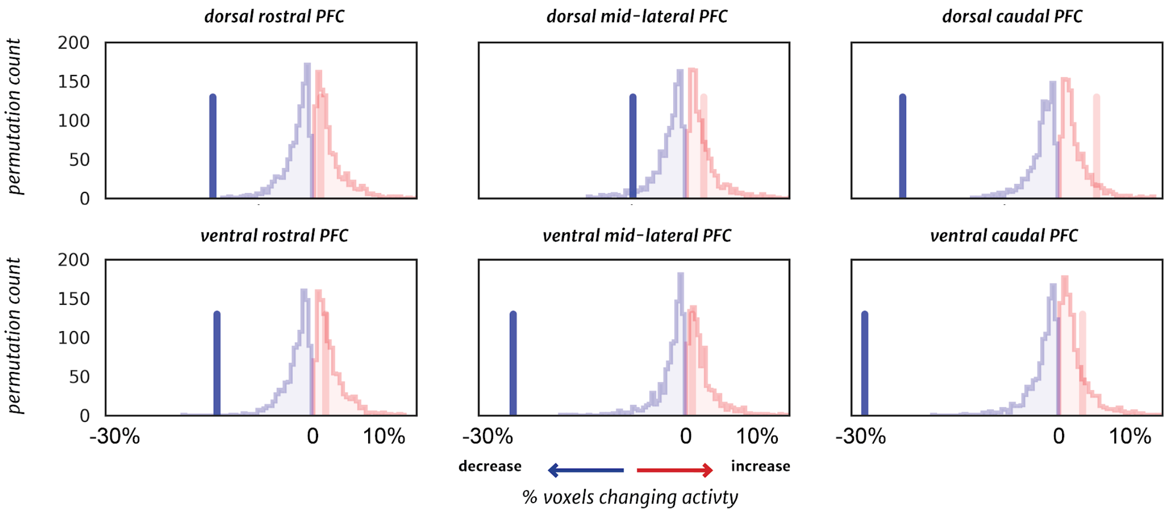


**SI Figure 3. Distribution of activity for the WM encoding epoch in PFC across the course of learning.** Significant increases (*red*) or decreases (*blue*) in the percentage of voxels changing activity across training are indicated by bolded vertical lines. Null distributions were created exactly as in Figure 3, but instead using the WM encoding period activity across sessions. All ROIs show a significant proportion of voxels with a decrease in activity, with no ROIs showing an increase in WM encoding activity across training.

### Individual sequence identity analysis

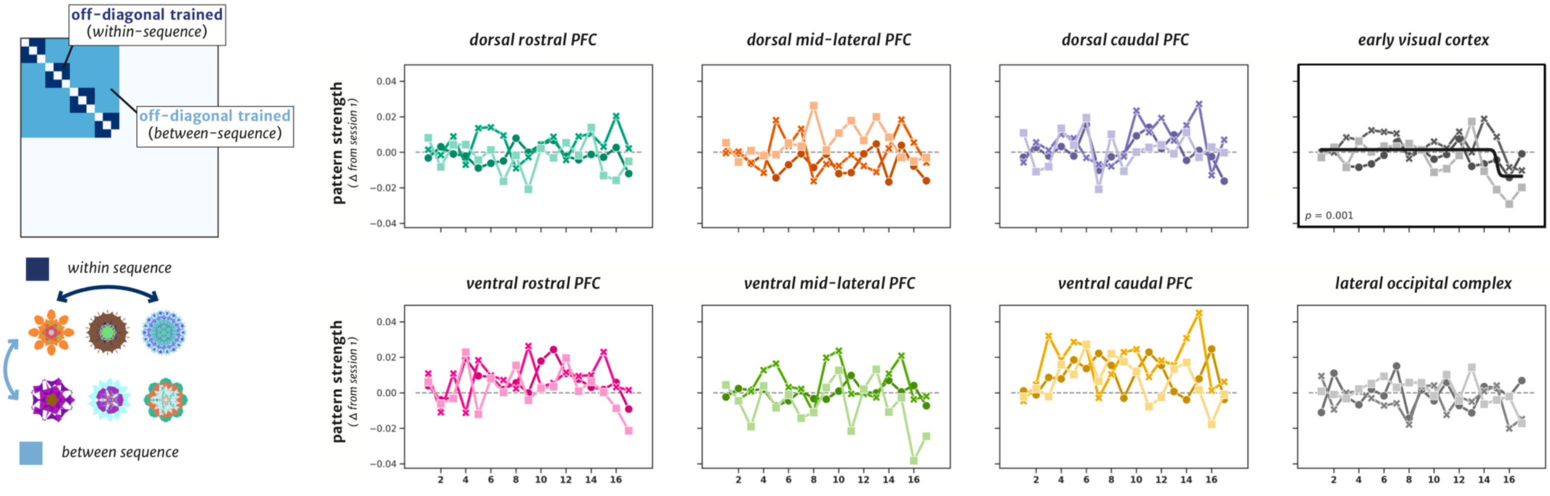


**SI Figure 4. Analysis of individual sequence representation in WM delay activity.**
**(a)** Left: Schematic of the model matrix for the analysis of correlations for items within the same trained sequences (dark blue, positive values) compared to correlations of items between different sequences (light blue, negative values). Right: Plots of the pattern strength across sessions for each ROI, as assessed by the model fit for the *individual sequence-level* model on the left. For visualization, all ROIs with significant changes in pattern strength across sessions are indicated with a *p*-value and bolded plot border, and pattern strength is plotted as a change from initial baseline values (session 1 and 2 mean). Only *early visual* cortex showed any significant change in model strength over time ($p$ = 0.001 [FDR-corrected p=0.004]; all other $p$-values > 0.2), and this was in the opposite direction than expected, with greater similarity over time for stimuli across different sequences. Each color shade represents one of the three individual participants.

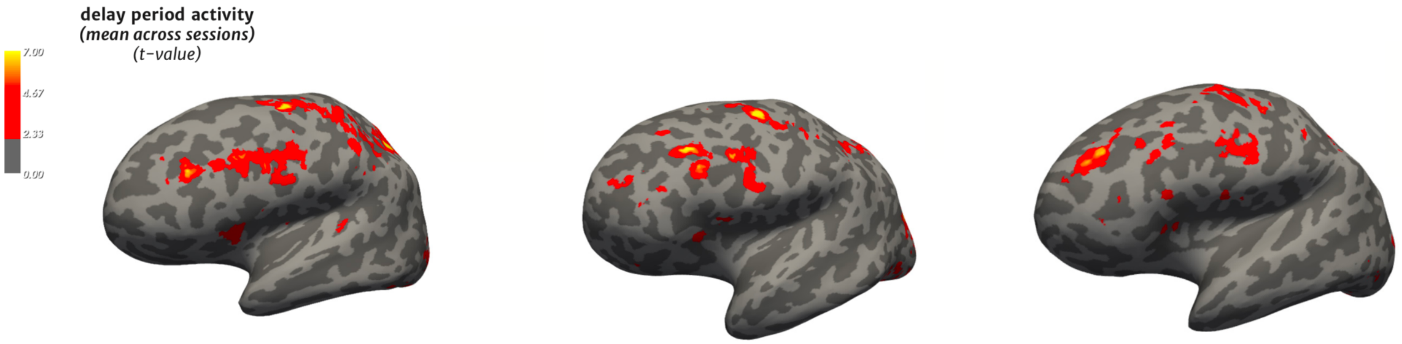


**SI Figure 5.**
Mean WM delay period activation ($t$-value, delay > fixation, thresholded $t > 2.5$) from each participant, averaged across all sessions. Light smoothing (1 step) was applied across the cortical surface for visualization. The unthresholded delay period activation maps from each of the 17 sessions are also available for viewing and download on the NeuroVault platform (https://identifiers.org/neurovault.collection:12687).

**Highly active** voxels ($t$ > 2.5 threshold)

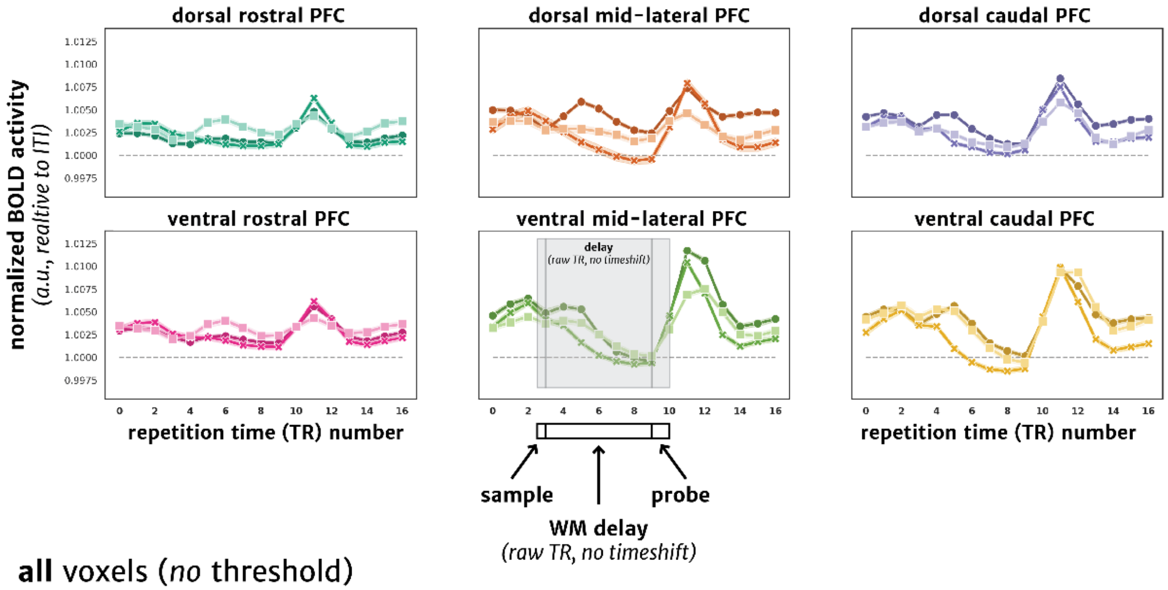


**all** voxels (*no* threshold)

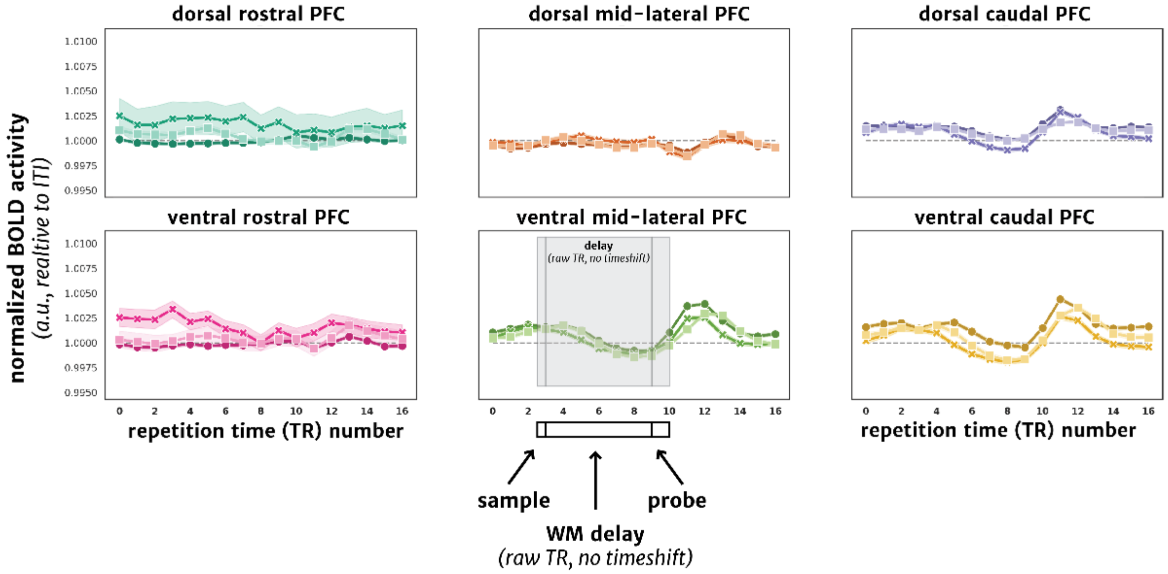


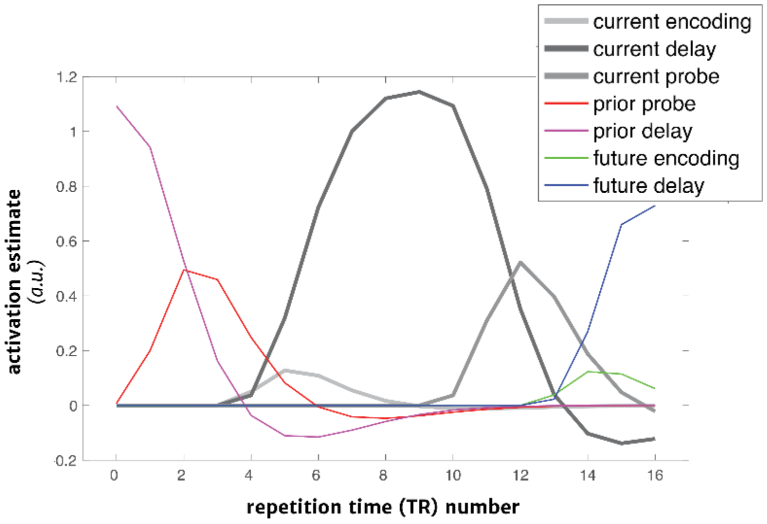

**SI Figure 6.**

Mean normalized BOLD activity (normalized, relative to ITI mean) for each of the six PFC and two visual regions across the time series of trials with a long (12 s) WM delay period for highly active (Top; $t > 2.5$) and all voxels (Middle). The beginning and end of the delay period are indicated with arrows and gray shading in the bottom center plot. Time points are raw and nto time-shifted to account for hemodynamic lag in the HRF. Line colors represent each of the three individual participants. Bottom: example time series of the predicted activation for trials as used to model the delay period activity. Different task phases are indicated in the legend for the encoding (0.5 s), delay (12 s), and probe (2 s) periods (current indicates trial beginning just after TR= 2, prior indicates preceding trial events, future indicates subsequent trial events).

**SI Results.**

Below we report the results for each RSA model from the ROIs highlighted in each Figures that showed significant fits using a logistic model. To ensure that our prior results were not dependent on fitting non-linear models to the data, we confirmed that reliable changes over time were found using a 2nd order model. To do this, the linear term of an ordinary least squares (OLS) model was analyzed to see if changes were also significant when modeled with a linear increase.

For the *item-level* model (**Figure 4b**), ventral mid-lateral PFC showed a significant linear increase in pattern strength across sessions ($\beta = 0.0004$, $t = 2.59$, $p = 0.013$).

In the c*ategory-level* model (**Figure 4c**), dorsal caudal PFC ($\beta = 0.0007$, $t = 2.56$, $p = 0.014$) and early visual cortex ($\beta = 0.0008$, $t = 2.89$, $p = 0.006$) also showed a significant linear increase in pattern strength across sessions, with ventral caudal PFC showing a marginal trend ($\beta = 0.0006$, $t = 1.96$, $p = 0.056$). However, in dorsal rostral PFC and ventral mid-lateral PFC, the logistic model fits were not apparent when testing for a linear increase ($\beta = 0.0004$, $t = 1.4$, $p = 0.16$; $\beta = 0.0002$, $t = 0.94$, $p = 0.35$).

Finally, in the *sequence category* model (**Figure 5**), both dorsal caudal PFC and ventral caudal PFC increases in pattern strength with the logistic model were replicated when testing for a significant linear increase across sessions (*dorsal caudal*: $\beta = 0.0002$, $t = 4.03$, $p < 0.001$; *ventral caudal:* $\beta = 0.0002$, $t = 2.93$, $p = 0.005$).

…

\
